# Cumulus: a cloud-based data analysis framework for large-scale single-cell and single-nucleus RNA-seq

**DOI:** 10.1101/823682

**Authors:** Bo Li, Joshua Gould, Yiming Yang, Siranush Sarkizova, Marcin Tabaka, Orr Ashenberg, Yanay Rosen, Michal Slyper, Monika S Kowalczyk, Alexandra-Chloé Villani, Timothy Tickle, Nir Hacohen, Orit Rozenblatt-Rosen, Aviv Regev

## Abstract

Massively parallel single-cell and single-nucleus RNA-seq (sc/snRNA-seq) have opened the way to systematic tissue atlases in health and disease, but as the scale of data generation is growing, so does the need for computational pipelines for scaled analysis. Here, we developed Cumulus, a cloud-based framework for analyzing large scale sc/snRNA-seq datasets. Cumulus combines the power of cloud computing with improvements in algorithm implementations to achieve high scalability, low cost, user-friendliness, and integrated support for a comprehensive set of features. We benchmark Cumulus on the Human Cell Atlas Census of Immune Cells dataset of bone marrow cells and show that it substantially improves efficiency over conventional frameworks, while maintaining or improving the quality of results, enabling large-scale studies.

---

Single-cell and single-nucleus RNA-seq (sc/snRNA-seq) revolutionized our ability to study complex and heterogeneous tissues, opening the way to charting cell atlases of complex tissues in health and disease, including the Human Cell Atlas^1^ and related initiatives. Advances in massively parallel sc/snRNA-seq^2,3^, now allow routine profiling of millions of cells^4,5^. Such large and growing datasets, however, pose a significant challenge for current analysis tools, which were designed to run on a local computer server and lack the computation capabilities required for processing terabytes of sequencing data.

To address this pressing challenge, we developed Cumulus, a cloud-based data analysis framework that is scalable, cost-effective, able to process a variety of data types and easily accessible to biologists (**Fig. 1**). Cumulus consists of a cloud analysis workflow, a Python analysis package (Pegasus), and a visualization application (Cirrocumulus). Cumulus performs three major steps in sc/snRNA-seq data analysis (**Fig. 1a**): (**1**) sequence read extraction; (**2**) gene-count matrix generation; and (**3**) biological analyses. It addresses them for big sc/snRNA-seq data by combining the power of cloud computing, algorithmic improvement, and more efficient implementation, as we describe below. To test Cumulus and compare it to other tools we relied on a scRNA-seq dataset of 274,182 cells (**Methods**), which were profiled from the bone marrow of 8 donors as part of the Human Cell Atlas Census of Immune Cells dataset^6^.

**Figure 1.**
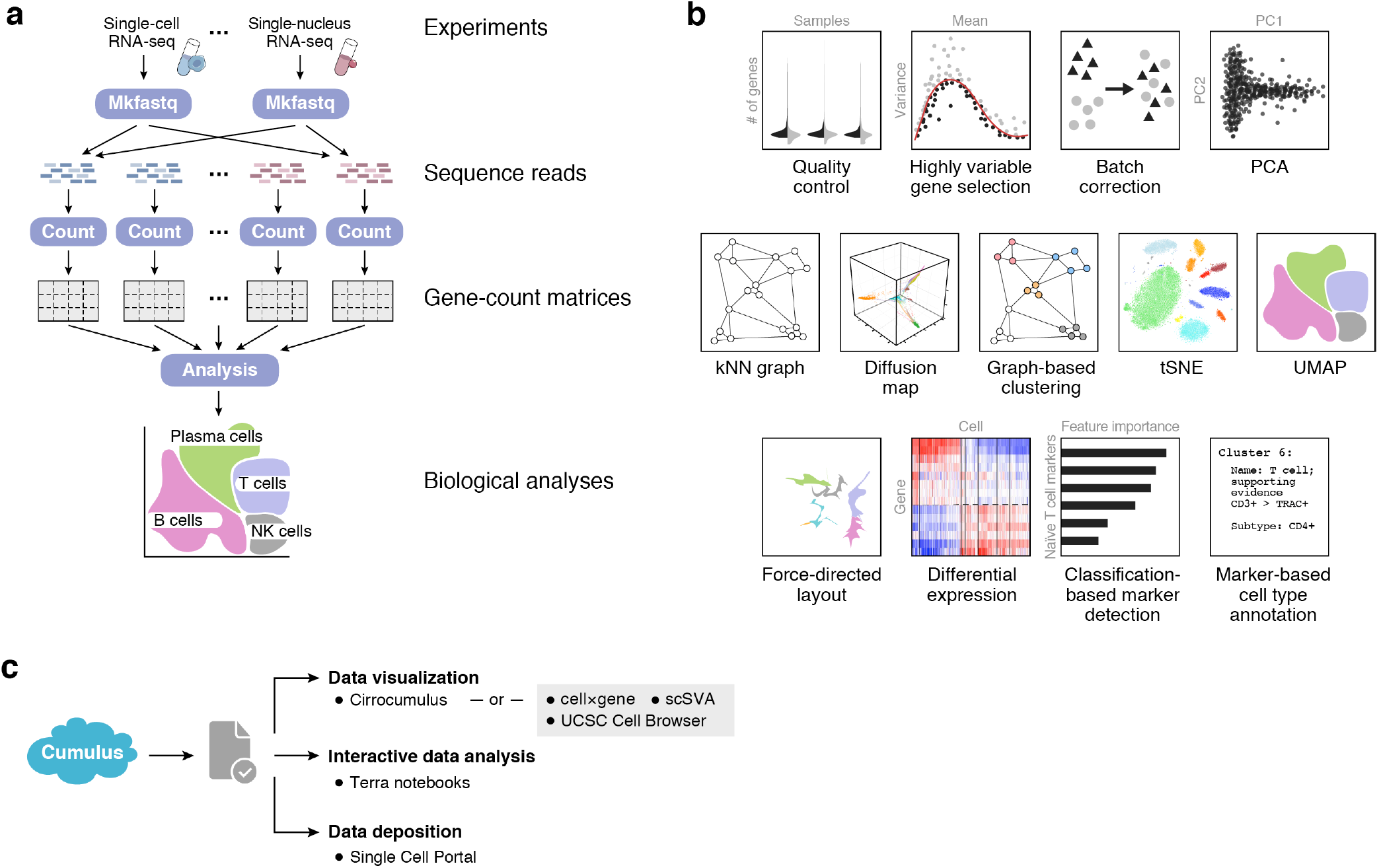
Cumulus: a scalable, feature-rich, accessible cloud based framework for sc/sn RNA-seq analysis. **a.** Cumulus data analysis workflow. Cumulus takes raw BCL files as input and outputs diverse analysis results, with three key computational steps – *mkfastq*, *count*, and *analysis*. **b.** sc/snRNA-seq analysis tasks in Pegasus. **c.** Cumulus enables flexible interactive data visualization and analysis. Users can instantly visualize Cumulus analysis results with Cirrocumulus, or publicly available visualization tools such as cellxgene, UCSC cell browser and scSVA. They can also interactively explore them on Terra Jupyter notebooks using Pegasus and deposit their data into the Single Cell Portal.

Cumulus leverages cloud computing and compatible data platforms. It is currently based on the Terra platform [https://app.terra.bio/] and Google Cloud Platform, but is generally cloud agnostic, as it depends only on Dockers and Workflow Description Language (WDL, **Methods**). Cumulus executes the first two steps – sequence read extraction and gene-count matrix generation – parallelly across a large number of computer nodes, and executes the last step of analysis in a single multi-CPU node, using its highly efficient analysis module (**Methods**, below). Cloud computing offers on-demand scalable computing, high-availability storage, data security and installation-free Software-as-a-Service (SaaS) capabilities, all at a low price. Non-programming biologist users readily access computing resources on the cloud through a simple web-based user interface provided by Terra (**Supplementary Video 1**).

Cumulus supports analysis starting from a variety of input modalities, such that scientists can use it as a single framework for diverse data types, all of which share a single cell/nucleus transcriptome as a core readout (**Supplementary Table 1**). These include: droplet-based^2,7^ (3’ or 5’ ends, with UMIs) and plate-based^8^ (full length, no UMI) sc/snRNA-seq (**Methods**); CITE-seq^9^, which simultaneously measures mRNA expression and the abundance of oligo-tagged surface antibodies in single cells (**Methods**), data from both cell^10^ or nucleus^11^ hashing experiments, which are lab techniques that reduce batch effects and cell/nucleus profiling costs, using a probabilistic demultiplexing algorithm^11^ (**Methods**); and Perturb-seq methods for pooled CRISPR screens^12–16^ with scRNA-seq readout (**Methods**). Other mainstream, non-Cloud based analysis packages, such as Cell Ranger^7^, Seurat^17^, or SCANPY^18^, currently support only some of these input data types, posing a potential burden for users (**Supplementary Table 1**).

The Cumulus analysis module, Pegasus, which can also run on the cloud or as an independent Python package, supports a comprehensive set of features, spanning most commonly used scRNA-seq analysis tasks (**Fig. 1b**, **Methods**). Starting from a gene-count matrix, Pegasus filters out low-quality cells/nuclei, selects highly variable genes (HVG) and optionally corrects batch effects. It then performs dimensionality reduction by principal component analysis (PCA) on HVGs, constructs a *k* nearest neighbor (*k*-NN) graph on the Principal Component (PC) space, calculates diffusion maps^19,20^ and applies community detection algorithms on the graph to find clusters^21,22^. It visualizes cell profiles using either t-SNE^23,24^-based or UMAP^25,26^-based methods. It can additionally estimate diffusion pseudotime^20^ and visualize developmental trajectories using force-directed layout embedding (FLE)^27–29^ based algorithms. Pegasus can be used to detect cluster-specific markers, by differential expression analysis between cells within and outside of a cluster and optionally calculates the area under ROC curve (AUROC) values for all genes (**Methods**). It can also train a gradient boosting tree classifier^30^ on the gene expression matrix to predict cluster labels and output genes with high feature importance scores (**Methods**), which provide additional information for detecting cluster-specific markers. Lastly, it annotates clusters with putative cell type labels based on user-provided gene sets (**Methods**). Pegasus thus offers diverse features, comparable to two other mainstream packages, Seurat^17^ and SCANPY^18^, although each package also has some unique features, absent from the other two (**Supplementary Table 2**).

Once the data are analyzed, users can visualize their results instantly using Cirrocumulus, a serverless application that enables interactive data visualization and sharing (**Fig. 1c, Supplementary Video 2**). Since Cirrocumulus only downloads to the browser those data that are necessary for visualization (**Methods**), it is scalable to millions of cells. Users can also download Cumulus-produced HDF5 result files for use with other visualization tools such as cellxgene^31^, UCSC Cell Browser^32^, and scSVA^29^ (**Fig. 1c**). Alternatively, users can inspect or re-analyze their data interactively using Pegasus on Terra Jupyter Notebooks (**Fig. 1c**). To help users better navigate data, we developed scPlot (**Methods**), a python package for generating interactive figures, as part of Pegasus. Finally, users can synchronize Cumulus results to the Single Cell Portal^33^ for data deposition (**Fig. 1c**). Cumulus is demoed as a featured workspace on Terra [https://app.terra.bio/#workspaces/fc-product-demo/scRNA-seq-cloud].

To ensure scalability, we enhanced the performance of the Pegasus analysis module through several algorithmic and implementation improvements in some of the most intensive tasks: the selection of highly variable genes (HVGs), batch correction, *k*-NN graph construction, calculation of diffusion pseuodotime (DPT), a combination of spectral and community-based clustering, and efficient visualization algorithms. We describe each of these enhancements in turn, comparing its impact on analysis quality and analysis speed/scale with other major packages.

First, we implemented a new HVG selection procedure that simplifies the calculation process and provides a mathematically sound way to handle batch effects (**Supplementary Fig. 1a, Methods**). For users’ convenience, we also include a standard procedure^17^, which is used by both SCANPY and Seurat. Comparing the new and standard procedures when applied to the bone marrow dataset suggests that the new procedure has at least equal quality *vs*. the standard one. It recovers slightly more immune-specific genes provided by the ImmPort^34^ data repository (**Supplementary Fig. 1b, Supplementary Data 1, Methods**), including important T cell markers, such as CD3D, CD3E and CD4, which are missed by the standard procedure. It also identifies one more cell type, megakaryocytes, which is missed by the standard procedure (**Supplementary Fig. 1c**).

Next, we enhanced the scalability of batch correction in Pegasus, by implementing the classical location and scale (L/S) adjustment method^35^, which relies only on linear operations and is thus much faster. As a benchmark, we ran Pegasus’s L/S method, SCANPY’s offerings of Combat^36^, MNN^37^ and BBKNN^38^, and the most recent integration method^39^ of Seurat v3 on a subset of the bone marrow dataset (**Methods**) and compared each method’s batch correction efficiency using two measurements, the kBET^40^ and kSIM acceptance rates. Briefly, the kBET acceptance rate measures if batches are well-mixed in the local neighborhood of each cell; the kSIM acceptance rate measures if cells of the same pre-annotated cell type are still close to each other in the local neighborhoods after batch correction (**Methods**), and helps reflect if known biological relations are preserved after correction. An ideal batch correction method should have both high kBET and kSIM acceptance rates. Each of the five methods evaluated showed a trade-off between the two rates, and none was a clear best performer (**Fig. 2a**), Pegasus’s L/S method is the fastest (**Supplementary Fig. 2a**), while maintaining a good balance between the two rates (**Fig. 2a, Supplementary Fig. 2b-g**).

**Figure 2.**
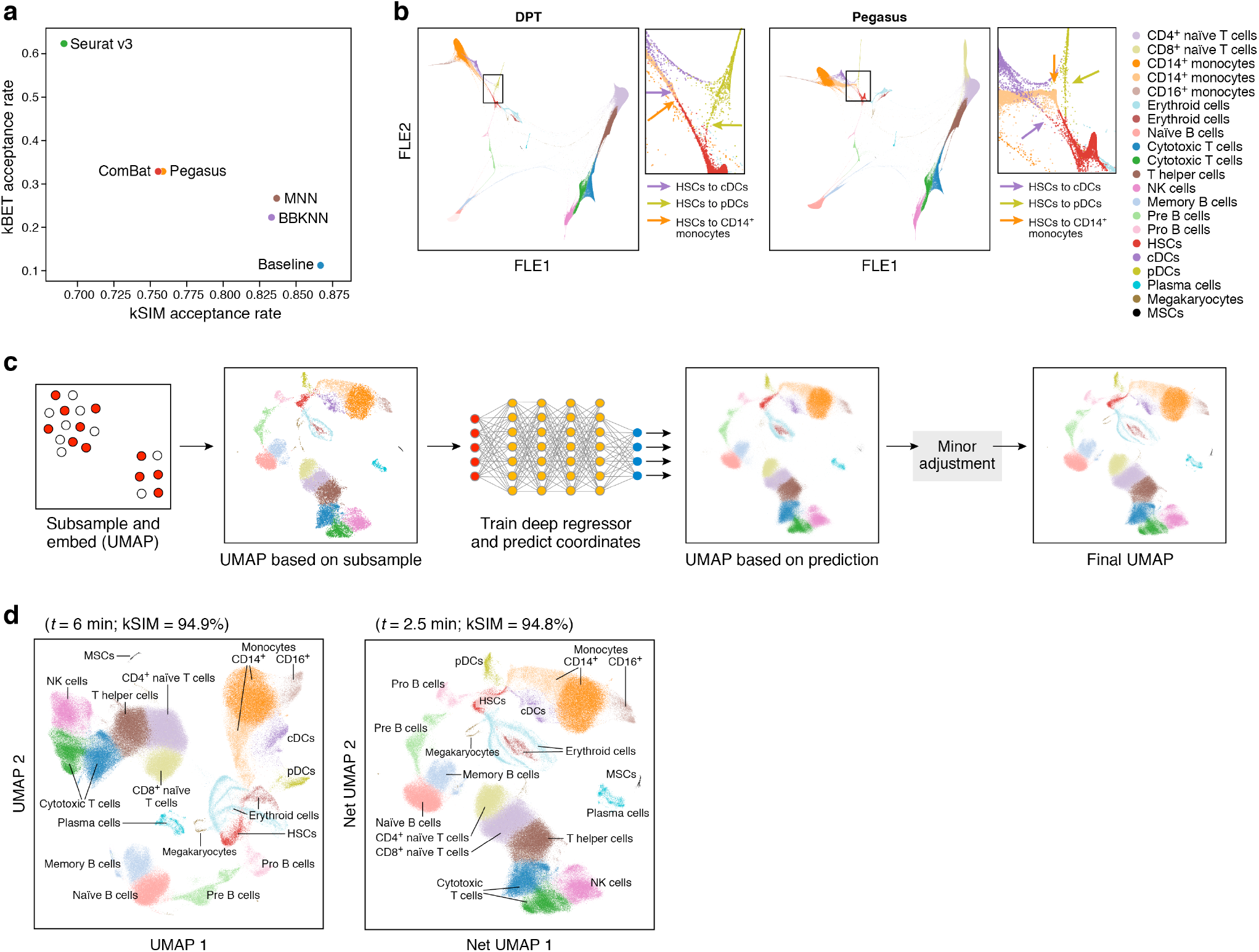
Algorithmic and implementation improvements underlying Pegasus’s high scalability. **a.** Trade-off between kBET and kSIM acceptance rates across different methods. kBET (*y* axis) and kSIM (*x* axis) acceptance rates of Pegasus, ComBat, MNN, BBKNN and Seurat v3 on 34,654 bone marrow cells. **b.** Improved resolution of a developmental bifurcation with diffusion pseudotime map with timescale selected by von Neumann entropy. Diffusion maps of cells colored by subset annotation (color legend), generated by DPT (left) and Pegasus (right). Red square: area of bifurcation from hematopoietic stem cells (HSCs) to CD14^+^ monocytes (orange arrow) and conventional dendritic cells (cDCs, purple arrow) (zoom, right), in each map. **c.** Deep-learning-based efficient visualization with Net-*. From left: a small fraction of cells is subsampled based on local density and then embedded (*e.g*., with UMAP); a deep regressor is trained on the subsampled cells to predict the embedding coordinates; it is then used to predict embedding coordinates for remaining cells; all the coordinates are fine-tuned by applying the embedding algorithm (*e.g*., UMAP) for a small number of iterations. **d.** Net-UMAP visualization is faster than UMAP while maintaining visualization quality. Embedding generated by UMAP (left) and Net-UMAP (right) of cells, colored by subset annotation. Top: Execution time and kSIM acceptance rate.

We also enhanced the scalability of *k*-NN graph construction by adopting the Hierarchical Navigable Small World (HNSW)^41^ algorithm, a state-of-the-art approximate nearest-neighbor finding algorithm, which was previously shown to be fastest for high quality approximations^42^. We compared HNSW with the approximate nearest neighbor finding algorithms used by SCANPY and Seurat on the bone marrow dataset based on speed and on recall, defined as the percentage of nearest neighbors that are also found by the brute-force algorithm (**Methods**). HNSW has a near optimal recall (**Supplementary Fig. 3a**), while being 3-19x faster (**Supplementary Fig. 3b**). HNSW was also benchmarked recently for plate based, small scale, scRNA-seq^43^.

To speed up the calculation of diffusion maps^19^ and diffusion pseudotime^20^ (DPT), we adopted two modifications, also used by SCANPY and scSVA, and further improved the identification of pseudotemporal trajectories^20^ with two additional modifications. As in SCANPY, we construct the affinity matrix based on the approximate *k*-NN graph constructed in the previous step instead of a complete graph, and also use only the top *n* diffusion components to approximate diffusion distances and thus diffusion pseudotimes, where *n* is a user-specific parameter. In addition, to better identify pseudotemporal trajectories when there are multiple subsets of cells undergoing separate temporal processes, we found that using more diffusion components helps us better separate different cell populations (**Supplementary Fig. 4a**, red regions) and thus we set *n* = 100 by default. We further introduce a family of diffusion pseudotime maps parameterized by timescale *t*: each pseudotime “meta-map” is constructed by summing over diffusion maps up to its timescale *t* (**Methods**). The DPT method^20^ is equivalent to the special case of this family with *t* = ∞. As timescale *t* increases, diffusion maps begin to smooth out local noise^44^. However, when *t* becomes too large, diffusion maps will also smooth out real signals^44^. Thus, instead of *t* = ∞, we choose a timescale *t* that smooths out most noise but little signal. Inspired by the PHATE^45^ method, we propose to pick *t* as the knee point on the curve of von Neumann entropies^46^ induced by diffusion maps at different timescales (**Supplementary Fig. 4b**, **Methods**). In the Immune Cell Atlas data, using the selected *t*, we identify a trajectory (**Fig. 2b**) that more clearly bifurcates from hematopoietic stem cells into CD14^+^ monocytes and conventional dendritic cells (cDCs), whereas those two lineages overlap in the DPT model.

In addition to offering popular modularity-based community detection algorithms for clustering cell profiles, including Louvain^21^ and Leiden^22^ (**Methods**), Pegasus also includes spectral-community-detection algorithms, such as spectral-Louvain and spectral-Leiden, which combine the strengths of both spectral clustering^47^ and community detection algorithms (**Methods**). Spectral clustering performed by applying the *k*-means algorithm on the calculated diffusion pseudotime components is very fast, but its clustering results are not always satisfactory (**Supplementary Fig. 5a**). Conversely, in Pegasus’s spectral clustering, we first aggregate cells into small groups and then apply community detection algorithms on the aggregated groups (**Methods**). On the bone marrow dataset, this new method provides clusters that are comparable to those from modularity-based community detection algorithms, but at the high speed of spectral clustering (**Supplementary Fig. 5b,c**).

Finally, in addition to visualization of single cell profiles using either t-SNE^23^, FIt-SNE^24^, UMAP^25^ (**Methods**), or a force directed layout embedding (FLE^28^) of the diffusion pseudotime map (**Methods**), we also include a deep-learning-based visualization technique that speeds up a generalized set of these and similar visualization algorithms (**Methods**). Inspired by net-SNE^48^, this technique is based on the assumption that large datasets are often redundant and their global structure can be captured using only a portion of the data. It thus first subsamples a fraction of cells according to each cell’s local density, ensuring higher rate of sampling from rare and sparse clusters, and then embeds the subsampled cells using the embedding algorithm of interest, such as UMAP (**Fig. 2c**). It then trains a deep-learning-based regressor (**Methods**) on the subsampled cells using their embedded coordinates as ground truth and uses the regressor to predict embedding coordinates for the remaining cells. Because the predicted coordinates yield a “blurry” visualization (**Fig. 2c**), it has a final refinement step, which applies the embedding algorithm for a small number of iterations to both the calculated coordinates for the subsampled cells and predicted coordinates for the remaining cells (**Fig. 2c**). We call visualizations obtained using this technique as Net-* visualizations and show that they speed up the original embedding algorithm by at least 2x and maintain the visualization quality based on similar kSIM acceptance rates (**Fig. 2d** and **Supplementary Fig. 6**).

As a result of these combined algorithmic and implementation improvements, Pegasus is much faster than other packages for running key analyses tasks on the bone marrow dataset^6^ (**Supplementary Table 3**, **Methods**) and the 1.3 million mouse brain dataset^4^ (**Supplementary Table 4**, **Methods**). In addition, with its cloud-based architecture, Cumulus is much faster than other packages when benchmarking on the bone marrow dataset. Compared to a Cell Ranger + Seurat/SCANPY pipeline, Cumulus completed the analysis in around 15 hours, while the alternative pipeline took over 9 days to run (**Table 1**, **Methods**). The associated computational costs were modest (*e.g*., ~$2 on average for around 4,000 cells in one sample, **Table 1**, **Methods**).

**Table 1.** Cumulus is computationally efficient and cost effective. Left columns: Total execution time on the bone marrow dataset of Cumulus, Cell Ranger + Seurat v3 or Cell Ranger + SCANPY pipeline, running on a 32 CPU thread, 120 GB memory Google Cloud virtual machine instance (**Methods**). Right columns: Average computational cost for running Cumulus per sample of ~4,000 cells (**Methods**).

|  | Time |  |  | Cost per sample |
| --- | --- | --- | --- | --- |
| Step | Cell Ranger + Seurat v3 | Cell Ranger + SCANPY | Cumulus | Cumulus |
| **Total** | **10 d, 5 h, 38 min** | **9 d, 5 h, 35 min** | **15 h, 15 min** | **\$1.832** |
| Mkfastq | 13 h, 18 min | 13 h, 18 min | 7 h, 54 min | \$0.22 |
| Count | 8 d, 14 h, 12 min | 8 d, 14 h, 12 min | 6 h, 44 min | \$1.61 |
| Analysis | 26 h, 8 min | 2 h, 5 min | 37 min | \$0.002 |

In conclusion, Cumulus provides the community with a cloud-based, scalable, cost-effective, comprehensive and easy to use platform for single-cell and single-nucleus RNA-seq research. Pegasus, Cumulus’ analysis module, which can also be used as an independent Python package, implements many improvements which enhance efficiency, from a new HVG selection procedure to a generalized deep-learning-based visualization speedup. While a complex framework such as Cumulus cannot provide an optimal combination of specialized algorithms for each application, it will accelerate research by providing an integrated and fast approach that enables more labs to analyze large-scale single-cell and single-nucleus datasets. In a separate study^49^, we demonstrated Cumulus by analyzing single-cell and single-nucleus RNA-seq data from fresh and frozen tumor from the Human Tumor Atlas Pilot Project (HTAPP). As the community produces data sets at substantially larger scales, we hope that Cumulus will play a key role in the effort to build atlases of complex tissues and organs at higher cellular resolution and leveraging them to understand the human body in health and disease.

## Code availability

Cumulus code consists of four components: the Pegasus and scPlot python packages, the Cumulus WDL workflows and Dockerfiles, the Cumulus docker images and the Cirrocumulus app. Pegasus source code is available at https://github.com/klarman-cell-observatory/pegasus. Pegasus_documentation is available at https://pegasus.readthedocs.io. scPlot source code is available at https://github.com/klarman-cell-observatory/scPlot. We wrote all workflows using the Workflow Description Language (WDL, https://github.com/openwdl/wdl) and encapsulated all software packages into Docker images using Dockerfiles. Cumulus WDL and Dockerfiles are available at https://github.com/klarman-cell-observatory/cumulus. Cumulus Docker images are available at https://hub.docker.com/u/cumulusprod. For Terra users, we additionally deposit Cumulus workflows in the Broad Methods Repository https://portal.firecloud.org/?return=terra#methods and provide a step-by-step manual at https://cumulus-doc.readthedocs.io. Cirrocumulus source code is available at https://github.com/klarman-cell-observatory/cirrocumulus. Pegasus, scPlot, Cumulus WDL files and Docker files, and Cirrocumulus are licensed under a BSD 3-clause license. In addition, we documented licenses for Cumulus dependencies in **Supplementary Data 2**. Due to 3^rd^ party licensing requirements, we can only provide Cell Ranger dockers without bcl2fastq2 and users can build their private bcl2fastq2-containing Dockers by following instructions listed in Cumulus documentation.

## Data availability

The bone marrow dataset is available at https://data.humancellatlas.org/explore/projects/cc95ff89-2e68-4a08-a234-480eca21ce79. The 1.3 million mouse brain data set is available at https://support.10xgenomics.com/single-cell-gene-expression/datasets/1.3.0/1M_neurons.

## Acknowledgments

We thank Jennifer Rood for help with manuscript editing, Leslie Gaffney for help with figure preparation, Eric Banks and Anthony Philippakis for advice on creating Cumulus’ featured Terra workspace, Christine O’Day and Ellen Law for advice on licensing Cumulus, and Christine O’Day additionally for help on preparing **Supplementary Data 2**, Danielle Dionne, Julia Waldman, Jane Lee and Karthik Shekhar for their contribution in generating the preview data set and sharing it openly pre-publication; and Mariam Maarouf and Deniz Erdogan for transferring the Pegasus namespace on Read the Docs to us.

## Author Contributions

B.L. and A.R. conceived the study, designed experiments and devised analyses. B.L. developed computational methods. B.L. J.G., S.S. and Y.Y. implemented code. B.L., J.G., S.S., Y.Y., M.T., O.A and Y.O. conducted computational experiments. M.S., M.S.K. and A.V. helped interpret results from the Immune Cell Atlas data used in this manuscript; T.T. helped with Terra cloud related development; N.H., O.R.R. and A.R. supervised work. B.L., Y.Y. and A.R. wrote the paper with input from all the authors.

## Methods

### CUMULUS MODULES

#### Gene-count matrix generation for droplet-based scRNA-seq

Cumulus supports gene-count matrix generation for 10x Genomics V2 and V3 chemistry using Cell Ranger. Cumulus first demultiplexes Illumina base call files (BCLs) for each sequencing flowcell by running *mkfastq* steps parallelly in different computer nodes. Each *mkfastq* job calls ‘cellranger mkfastq’ to generate sequence reads in FASTQ files. By default, each *mkfastq* job requests 32 CPUs, 120 GB memory and 1.5 TB disk space from the cloud. Cumulus then generates gene-count matrices for each 10x channel by running *count* steps in parallel. Each *count* job calls ‘cellranger count’ with appropriate parameters and requests 32 CPUs, 120 GB memory and 500 GB disk space from the cloud by default. Cumulus also supports gene-count matrix generation for Drop-seq^2^ data using either the methods described in Drop-seq alignment cookbook^50^ or dropEst^51^.

#### Gene-count matrix generation for plate-based scRNA-seq

Cumulus supports gene-count matrix generation for scRNA-seq data generated by the SMART-seq2 protocol^8^ from sequence reads in FASTQ files. Cumulus estimates gene expression levels for each single cell in parallel in different computer nodes. Each node runs RSEM^52^ with default parameters and utilizes Bowtie 2^53^ to align reads. Each node requests 4 CPUs, 3.6 GB memory and 10GB disk space by default. Once expression levels are estimated, Cumulus converts the relative expression levels (in Transcript per 100K, TP100K) into a count vector for each single cell using the formula below and then generates a gene-count matrix by concatenating count vectors from all cells.

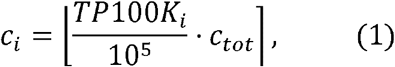

where *c_i_* and *TP*100*K_i_* are the converted read count and estimated expression level of gene *i*, respectively. *c_tot_* is the sum of RSEM-estimated expected counts from all genes.

#### Pegasus

Cumulus runs the *analysis* step on a single node, which requests 32 CPUs, 200 GB memory and 100 GB disk space by default. The *analysis* step calls Pegasus, a fast Python package we have implemented. Pegasus utilizes SCANPY’s AnnData data structure^18^ to store gene-count matrices and analysis results. More implementation details are discussed in the subsequent sections and **Supplementary Note 1**.

#### Feature-count matrix generation for CITE-seq, cell hashing, nucleus hashing, and Perturb-seq

Cumulus supports feature-count matrix generation of CITE-seq^9^, cell hashing^10^, nucleus^11^ hashing and Perturb-seq^12-16^ protocols, using either 10x Genomics V2 or V3 chemistry. Each feature-count matrix generation job runs parallelly on a separate compute node with 1 CPU, 32 GB memory and 100 GB disk space, and calls ‘generate_count_matrix_ADTs’, a fast C++ program we implemented, to extract the matrix from sequence reads in FASTQ files. The C++ program scans each read pair to search for valid sequence structures. We assume read 1 records the cellular barcode and Unique Molecular Identifier (UMI) information and read 2 records feature barcode information, such as hash tags for hashing protocols or sgRNA information for Perturb-seq (below). The first 16 nucleotides of read 1 represent the cell barcode for both V2 and V3 chemistry. The next 10 and 12 nucleotides represent the UMI for V2 and V3 chemistry, respectively. We allow up to 1 and 0 mismatch for matching cell barcodes in V2 and V3 chemistry, respectively.

Feature barcode information is recorded differently in read 2 for different protocols. For CITE-seq, cell hashing and nucleus hashing protocols, the location of the feature barcode depends on what type of BioLegend TotalSeq™ antibodies users choose. If TotalSeq™-A antibodies are used, the feature barcode is located at the 5’ end of read 2. Otherwise, the feature barcode starts at the 11^th^ nucleotide from the 5’ end of read 2. ‘generate_count_matrix_ADTs’ automatically detects antibody type by scanning read 2 of the first 1,000 read pairs and calculating the percentage of read pairs containing the auxiliary sequence. If more than 50% of read pairs contain the auxiliary sequence, we assume the antibody type is TotalSeq™-A, otherwise it is TotalSeq™-B or TotalSeq™-C. We allow up to 1 mismatch for matching the auxiliary sequence.

For Perturb-seq protocols, we assume that the feature barcode (protospacer) is located in front of a user-provided anchor sequence. For V2 chemistry, we first search the anchor sequence in read 2, allowing up to 2 mismatches or indels. We then extract the feature barcode at the 5’ end of the anchor sequence. For V3 chemistry, we assume users use 10x Genomics CRISPR guide capture assays and additionally check the Template Switching Oligo (TSO) sequence ‘AAGCAGTGGTATCAACGCAGAGTACATGGG’ at the 5’ end of read 2, allowing up to 3 mismatches and indels.

Once we locate the feature barcode, we match it with a user-provided white list, allowing up to 3 mismatches by default. After scanning all read pairs, ‘generate_count_matrix_ADTs’ generates a feature-count matrix in CSV format: each row represents one feature, each column represents one cell barcode, and each element records the number of unique UMIs for the feature in the row in the cell barcode in the column. To speed up sequence matching, we encode cell barcodes, UMIs and feature barcodes into 8-byte unsigned integers (2 bits per nucleotide).

#### CITE-seq data analysis

Based on the generated feature-count matrix, Cumulus first calculates the log fold change between feature UMI counts of the antibody of interest and its IgG control as the antibody expression, provided that users include both antibodies of interest and their corresponding IgG controls in their CITE-seq assays. Let us denote the UMI counts of the antibody and its IgG control as *c_a_* and *c_c_*. The antibody expression *expr* is calculated as

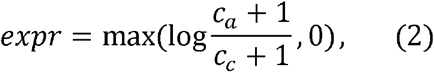

where we add 1 to both the numerator and denominator to avoid log 0. If IgG controls are not provided, we calculate *expr* as

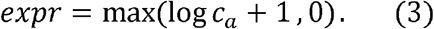

Cumulus merges the transformed antibody expression matrix into an RNA expression matrix so that users can plot antibody expression in 2D visualizations (*e.g.*, t-SNE & UMAP) calculated based on RNA expression levels. Cumulus can optionally generate t-SNE plots solely based on antibody expression levels.

#### Demultiplexing cell hashing and nucleus hashing data

Cumulus demultiplexes cell hashing and nucleus hashing data using the DemuxEM algorithm, which we recently described^11^.

#### Chimeric read filtration for Perturb-seq data

In Perturb-seq, sgRNAs are often amplified by dial-out PCR^12^ to ensure feature detection, and the resulting library is often over-sequenced, which can lead to a high number of false positive UMIs due to PCR chimeric reads^54^. Such false positive UMIs tend to have fewer supporting reads on average. Suppose we have c[i] UMIs with exact *i* supporting reads. In general, we expect *c*[*i*] to decrease monotonically as *i* increases. However, if the library is over-sequenced, we may observe a second peak in the tail of the *c*[*i*] distribution (∃*i, c*[*i* − 2] < *c*[*i* − 1] < *c*[*i*] ≥ *c*[*i* + 1]), which is more likely to represent true UMIs. Cumulus detects the left boundary of the second peak by scanning *i* consecutively. If Cumulus can find an *i* such that *c*[*i*] < *c*[*i* + 1] < *c*[*i* + 2] and *i* ≤ 10, Cumulus will filter out any UMIs with fewer than *i* supporting reads. Otherwise, Cumulus filters out any UMIs with only one supporting read. If a cell barcode and UMI combination contains more than 1 feature barcode, it is likely that the feature barcode with fewer supporting reads is produced by PCR chimeras^54^ and Cumulus will filter feature barcodes supported by no more than 10% of reads belonging to that combination. Cumulus generates a filtered feature-count matrix after this filtration step and lets users decide if they want to use the original feature-count matrix or the filtered feature-count matrix.

#### Cirrocumulus implementation

Cirrocumulus is a Google App Engine application for visualizing variables on a 2D or 3D embedding of observations. The client side of Cirrocumulus is implemented using React to manage state and Plotly to generate charts. The backend consists of several cloud functions to manage datasets stored in a NoSQL cloud database and to slice variables from a specified dataset in PARQUET format, where the PARQUET file is generated by Pegasus. The slice function can optionally generate statistical summaries on an n-dimensional grid, thus enabling plotting of millions of cells.

#### scPlot implementation

scPlot [https://github.com/klarman-cell-observatory/scPlot] is a plotting library included as part of Pegasus. Plots provided include scatter plots, feature plots, dot plots and violin plots and can scale to millions of cells by plotting cells on a two-dimensional grid. scPlot uses HoloViews [http://holoviews.org/], thus allowing the same code to generate interactive plots with Bokeh for a Jupyter notebook and static plots with Matplotlib.

#### Preprocessing

Pegasus selects high quality cells based on a combination of the following criteria, with user-provided parameters: 1) number of unique molecular identifiers (UMIs) between [*min_umis, max_umis*), default: *min_umis* = 100 and *max_umis* = 600,000; 2) number of expressed genes (at least one UMI) between [*min_genes, max_genes*), default: *min_genes* = 500 and *max_genes* = 6000; 3) percentage of UMIs from mitochondrial genes less than *percent_mito*, default: *percent_mito* = 10%. Pegasus then selects robust genes, defined as genes detected in at least *x* percentage of cells, where *x* is a user-defined parameter; default: *x* = 0.05% (equivalent to 3 cells out of 6,000 cells). Next, Pegasus normalizes the count vector of each cell, such that the sum of normalized counts from robust genes is equal to 100,000 transcripts per 100K (TP100K), and transforms the normalized expression matrix into the natural log space by replacing expression value *y* into log(*y* + 1). Additional details are available in **Supplementary Note 1**.

#### Highly variable gene selection

The standard HVG selection procedure operates in the original expression space. However, almost all downstream analyses are conducted in log expression space. To reconcile this inconsistency, we develop a new HVG selection procedure that operates directly in log expression space.

We select HVGs only from robust genes. Suppose we have *N* cells and *R* robust genes. We denote the log expression of gene *g* in cell *i* as *Y_ig_*. We first estimate the mean and variance for each robust gene *g* as

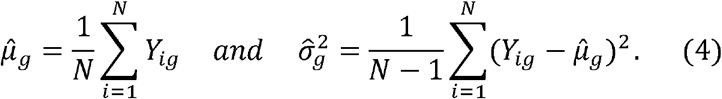

We then fit a LOESS^55^ curve of degree 2 (span parameter 0.02) between the estimated means and variances (**Supplementary Fig. 1a**) and denote the LOESS-predicted variance for gene *g* as 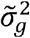. Any gene *g* with 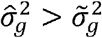 has a higher than expected variance.

We calculate the difference and fold change between the estimated and LOESS-predicted variances as

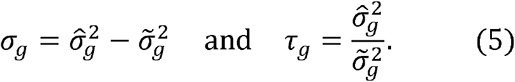

We then rank each robust gene with respect to *σ_g_* and *τ_g_* in descending order, and denote their rankings as *rank_σ_*(*g*) and *rank_τ_*(*g*) respectively. Lastly, we define the overall ranking as the sum of the two rankings

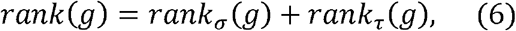

and select the top *n* robust genes with respect to *rank*(*g*) as HVGs.

The new procedure handles batch effects naturally. Suppose we have *K* biologically different groups, each group *k* has *n_k_* batches and each batch *kj* has *n_kj_* cells. We additionally denote the mean within batch *kj* and within group *k* as 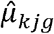 and 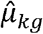, respectively. Because we have

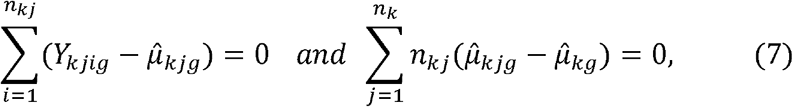

We can decompose the variance 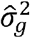 into three components – within-batch variance 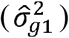, between-batch variance 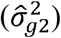 and between-group variance 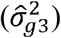 – as follows:

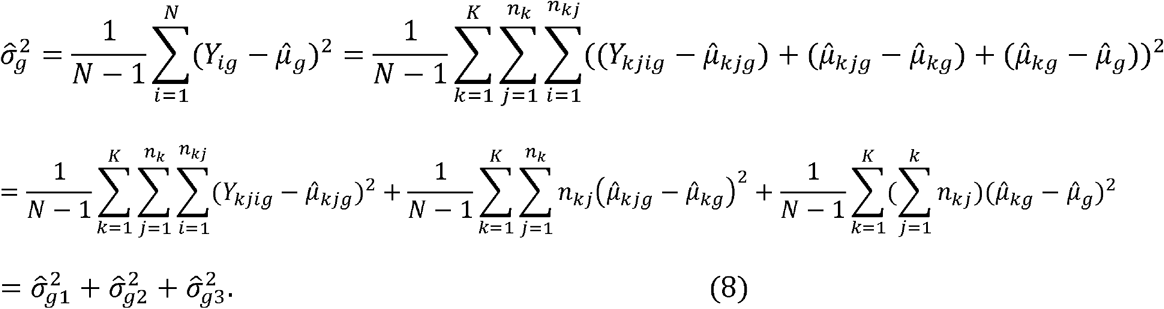

We remove the variance term 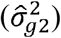 due to batch effects by redefining 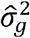 as

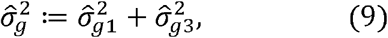

and plug in the new redefined variance term to the previously described procedure to select HVGs. If we use the redefined variance terms in HVG selection, we are in the batch-aware mode. We provide a detailed description of the new procedure in the **Supplementary Note 1 1**.

We also implemented the standard HVG selection procedure, which handles batch effects using the method in Seurat v3^39^, and documented implementation details in the **Supplementary Note 1**.

#### Batch correction with the L/S adjustment method

For simplicity, let us assume that we only have one biological group with *m* batches and each batch *j* has *n_j_* cells. We model the log gene expression level of gene *g* at batch *j*’s *i*th cell as

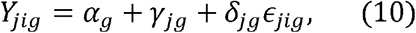

where *α_g_* is the baseline expression level of gene *g, ϵ_jig_* is the error term, which follows a distribution with a mean 0 and a variance 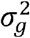. In addition, *γ_jg_* and *δ_jg_* are the additive and multiplicative batch effects, respectively. We estimate these parameters for each gene separately as follows:

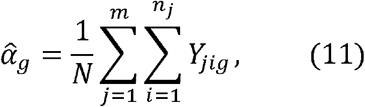

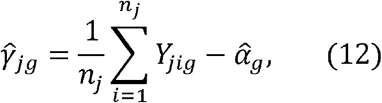

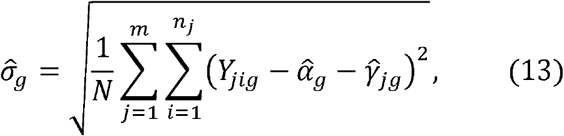

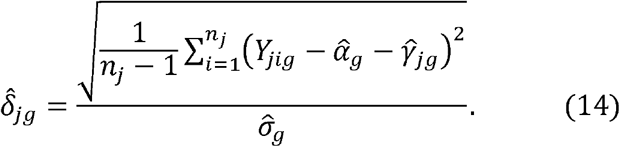

We denote 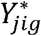 as the batch adjusted expression level, which is calculated as

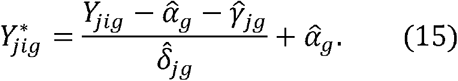

We provide a more detailed description of the L/S method in the **Supplementary Note 1**, including how to handle multiple biological groups.

Since batch correction transforms a sparse expression matrix into a dense matrix, which uses much more memory, we only calculate batch-adjusted expression levels for genes of interest, such as HVGs. We rewrite (15) as

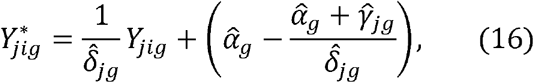

and use a two-step procedure to correct batch effects: First, we calculate and save batch-correction factors 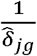 and 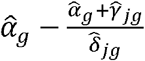 for all genes. Second, we calculate adjusted expression levels only for genes of interest using (16). We save the batch-correction factors for all genes, such that we can calculate batch-adjusted expression levels for any gene instantly in the future.

#### kBET acceptance rate

kBET^40^ acceptance rate measures if cells from different batches mix well in the local neighborhood of each cell. Pegasus implements the kBET acceptance rate calculation procedure as follows: We define *f* = (*f*_1_, …, *f_m_*) as the ideal batch mixing frequency, where 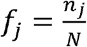. For each cell *i*, we find its *k* nearest neighbors (including itself) using the HNSW algorithm^41^ and denote the number of neighbors belonging to batch *j* as 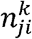. Then we calculate its *χ*^2^ test statistic with *m* − 1 degrees of freedom as

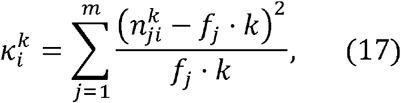

and its p value as

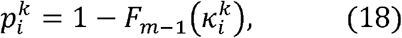

where *F*_*m*−1_(*x*) is the cumulative density function.

The kBET acceptance rate is calculated as the percentage of cells that accept the null hypothesis at significance level *α*:

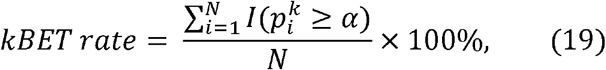

where *I*(*x*) is the indicator function, and *k* and *a* are user-specified parameters.

#### kSIM acceptance rate

The kSIM acceptance rate requires ground truth cell type information and measures if the neighbors of a cell has the same cell type as it does. If a method over-corrects the batch effects, it will have a low kSIM acceptance rate. We use the HNSW algorithm to find *k* nearest neighbors (including the cell itself) for each cell *i* and denote the number of neighbors that have the same cell type as *i* as 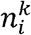. In addition, we require at least *β* fraction of neighbors of cell *i* to have the same cell type as *i* in order to say cell *i* has a consistent neighborhood. The kSIM acceptance rate is calculated as follows:

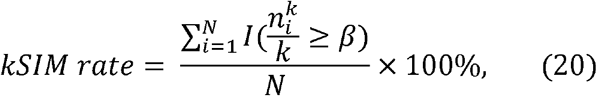

where *k* and *β* are user-specified parameters.

#### Dimensionality reduction by Principal Components Analysis

Pegasus calculates the top *m* principal components based on highly variable genes. It utilizes the randomized PCA algorithm^56^ implemented in *Scikit-learn* package^57^ to speed up the computation. By default, Pegasus sets *m* = 50.

#### *k*-nearest neighbors (*k*-NN) graph construction

Pegasus uses the HNSW^41^ algorithm with parameters *M* = 20, *efC* = 200, *efS* = 200, to construct k-NN graphs. By default, Pegasus searches the top 100 nearest neighbors (including the cell itself) for each cell (*K* = 100). Because HNSW is an approximate algorithm, it cannot always return the cell itself as the 1^st^ nearest neighbor. For any cell missing itself as the 1^st^ nearest neighbor, Pegasus sets itself as the 1^st^ nearest neighbor and picks the top 99 nearest neighbors returned by HNSW as the 2^nd^ to 100^th^ nearest neighbors. HNSW has a random index building process, which produces different indices in different runs if multiple threads are used. For reproducibility purposes, Pegasus provides two modes of running HNSW: robust mode and full speed mode. In robust mode, Pegasus runs the index building process with only one thread and runs the neighbor searching process with multiple threads. In full speed mode, Pegasus also runs the index building process with multiple threads. In either mode, Pegasus stores the neighbor searching results in the AnnData^18^ object. Without explicit notification, Pegasus runs HNSW in the robust mode.

#### Diffusion maps and diffusion pseudotime maps

We provide a high-level summary here and a more detailed description in **Supplementary Note 1**.

To compute diffusion maps, we first construct an affinity matrix **W**_*N×N*_ based on the top *m* principal components. This affinity matrix is also used in community-detection-based clustering algorithms. We construct **W** based on the top *K* nearest neighbors found by the HNSW algorithm. Let us define a cell **x**’s neighborhood set *N*(**x**) as the set consisting of **x**’s 2^nd^ to *K*^th^ nearest neighbors. We then define the following locally scaled Gaussian kernel between any two cells **x** and **y**:

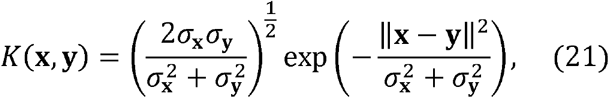

where, **x** is a vector containing the top *m* PC coordinates of cell **x**, and *σ*_x_ is **x**’s local kernel width, defined as *σ*_x_ = median{*d_i_*|*i*=2, …, *K*}, where *d_i_* is the distance between cell **x** and its *i*^th^ neighbor. To eliminate the effects of sampling density, we additionally define the following density-normalized kernel^19^:

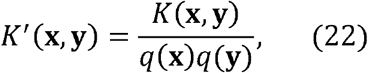

where *q*(**x**) is the sampling density term of cell **x** and defined as:

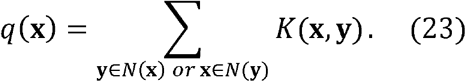

The affinity matrix **W** is constructed using the density-normalized kernel:

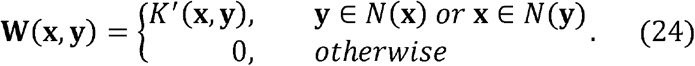

We then calculate the Markov chain transition matrix **P** and the symmetric “transition” matrix **Q** based on the affinity matrix:

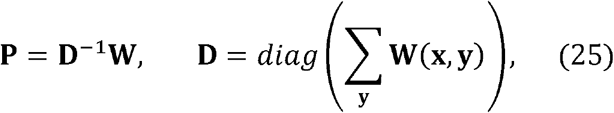

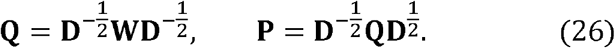

Since **Q** is symmetric, it has the eigen decomposition of **Q** = **UΛU**^*T*^. In addition, we know that in practice all **Q**’s eigenvalues are in (−1,1] and 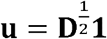 is its eigenvector for eigenvalue *λ* − 1 (**Supplementary Note 1**). We also know that **P** shares the same eigenvalues as **Q** and its right eigenvectors **Ψ** and left eigenvectors **Φ** are

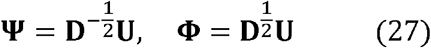

Next, to speed up the calculation, we approximate diffusion maps using only the top *n* diffusion components (**Supplementary Note 1**), where *n* is a user-specified parameter with default value *n* = 100. First, we calculate the top *n* eigenvalues and eigenvectors of **Q** using the Implicitly Restarted Lanczos Method^58^ (via scipy.sparse.linalg.eigsh function). We also provide the alternative option to calculate the top n eigenvalues using the randomized SVD algorithm^56^ (**Supplementary Note 1**). We order these *n* eigenvalues by magnitude:

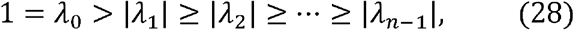

and define a family of approximated diffusion maps 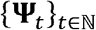 parameterized by timescale *t*:

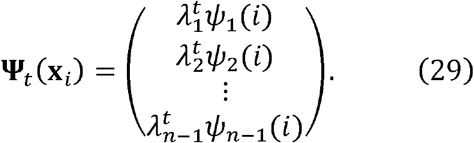

Note that we use the right eigenvectors of the transition matrix **P** to construct diffusion maps. Using the eigenvectors of **P** is consistent with the original diffusion map paper^19^ and recommended in the spectral clustering literature^47^. The DPT paper^20^ constructs diffusion maps using eigenvectors of the symmetric matrix **Q** instead.

We next define approximated diffusion pseudotime maps 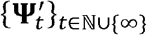 based on approximated diffusion maps:

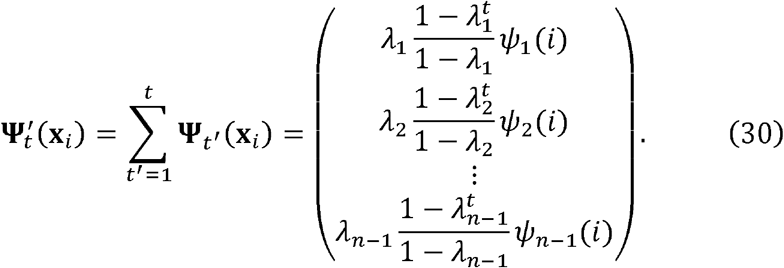

In particular, when *t* = ∞, we recover the DPT method (except it uses the eigenvectors of **Q**). We wish to pick a timescale *t* that smoothens out most of the noise but little signal. We select *t* based on the von Neumann entropy^46^ of the graph induced by each timescale. For each *t*, its power matrix **P**^*t*^ induces a graph with the following Laplacian and density matrices (**Supplementary Note 1**):

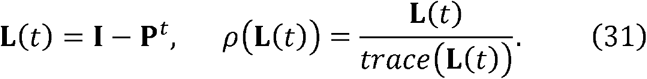

We derive the von Neumann entropy *S*(*t*) for the top *n* diffusion components from the density matrix *ρ*(**L**(*t*))(**Supplementary Note 1**):

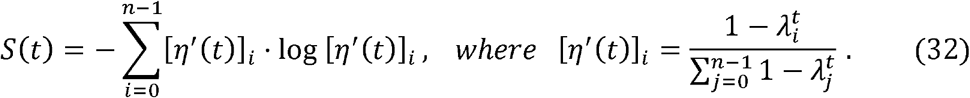

*S*(*t*) increases as *t* increases and reaches its maximum log (*n* − 1) when *t* → ∞. Because smaller eigenvalues of **P** (likely representing noise) decrease to 0 (and hence contribute a 1 to the entropy) much more rapidly than large eigenvalues (likely representing signal)^45^, we expect to observe a high rate of increase in *S*(*t*) initially when noise is smoothed out and then a low rate of increase when signal begins to be removed (**Supplementary Fig. 4b**). We pick the timescale *t* as the knee point of this von Neumann entropy curve.

Let us denote 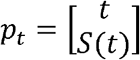 as the point for timescale *t* in the curve. We find the kneepoint by scanning all *t*s (where *t* is an integer) between 1 and *maxt*, and pick the *t* that is furthest from *p*_2_−*p*_1_, the segment connecting the two endpoints of the curve. *maxt* is a user-provided parameter set to *maxt* = 5000 by default. We calculate the distance between point *p_t_* and the segment *p*_2_ − *p*_1_ as follows:

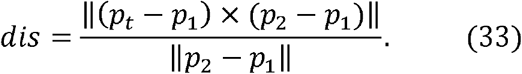

For the selected *t*, given a user-specified cell **r** as the root, we calculate the diffusion pseudotime distance from root to any other cell **x** as

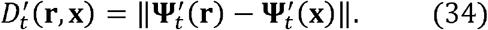

We then normalize the diffusion pseudotime distance into [0, 1] as the diffusion pseudotime.

#### Modularity-based community detection algorithms

We construct a weighted undirected graph *G* = (*V, E, w*) from the affinity matrix **W**. In the graph, vertex set *V* contains all cells, and an edge (*u, v*) ∈ *E* if and only if **W**_*u,v*_ > 0. The weight of the edge is calculated as:

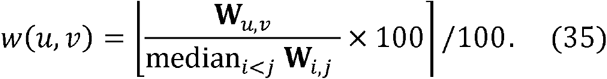

Community detection algorithms try to find a partition *C* of cells that maximizes the following modularity function^59^:

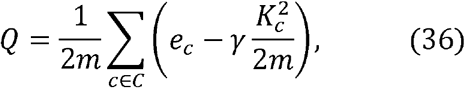

where each *c* ∈ *C* consists of cells in that community, *γ* is the resolution parameter controlling the total number of communities, and

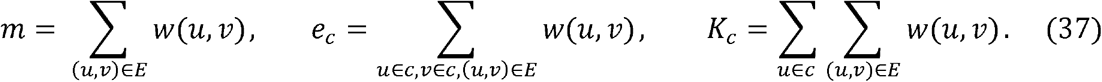

Pegasus supports two modularity-based community detection algorithms: Louvain^21^ and Leiden. For both algorithms, Pegasus sets the resolution *γ* = 1.3 by default and reports each community as a separate cluster.

The Louvain algorithm^21^ optimizes the modularity function *Q* in two phases: (1) in the move phase, each node is inspected and moved to the community that yields the largest increase in *Q*;(2) in the aggregation phase, each community aggregates into a new node to form an aggregated graph. The algorithm starts from the partition that each cell is its own community and repeats the two phases until there is no increase in *Q*. Pegasus uses the louvain-igraph implementation from Vincent Traag [https://github.com/vtraag/louvain-igraph]. Note that the latest release of louvain-igraph package (v0.6.1) contains a bug that prevents it from being reproducible even when the same random seed is used. Thus, Pegasus installs this package directly from the github master branch, in which the bug is fixed.

The Leiden algorithm^22^ is a recent improvement over the Louvain algorithm and consists of three phases: (1) a <u>move phase</u>, which is similar to Louvain’s; (2) a <u>refinement phase</u>, when each community found in (1) is examined and may be split into sub-communities; (3) an <u>aggregation</u> phase, when each sub-community from (2) is aggregated into a new node and assigned to an initial partition based on communities from (1). Pegasus uses the leidenalg implementation from Vincent Traag [https://github.com/vtraag/leidenalg]. Applying the Leiden algorithm on communities detected from previous Leiden runs can further improve the modularity function^22^. Thus, following SCANPY^18^, Pegasus runs the Leiden algorithm iteratively on the graph *G* until *Q* does not further improve (n_iterations = −1 by default).

#### Spectral-community-detection algorithms for fast clustering

Pegasus provides two spectral-community-detection algorithms: spectral-Louvain and spectral-Leiden. Spectral-community-detection algorithms aggregate cells into thousands of groups of cells, where each group consists of cells that are likely from the same “real” cluster, and then apply community detection algorithms such as Louvain and Leiden on the groups instead of on individual cells to achieve a major speedup.

Our variant of the spectral clustering algorithm partitions cells into groups by applying the *k*-means algorithm on calculated diffusion pseudotime component space, using a 2-level clustering strategy. We first partition the cells into *k*_1_ clusters using the scikit-learn’s KMeans function with default parameters. We further partition each of the *k*_1_ clusters into *k*_2_ sub-clusters using the KMeans function with *n_init* = 1. This procedure will give us *k*_1_ · *k*_2_ cell groups, on which we then apply community detection algorithms. Pegasus sets *k*_1_ = 30 and *k*_2_ = 50 by default. If diffusion pseudotime components are not calculated, Pegasus will apply the *k*-means algorithm on the PC space instead.

#### t-SNE, UMAP and FLE

Pegasus calculates a t-SNE using the Multicore-TSNE package implemented by Dmitry Ulyanov [https://github.com/DmitryUlyanov/Multicore-TSNE]. We found and fixed a random-seed-related bug in this package that prevents the package from reproducing the exact t-SNE coordinates, and provide a bug-free version of the package at [https://github.com/lilab-bcb/Multicore-TSNE]. Pegasus uses the following t-SNE parameters by default: perplexity=30, early_exaggeration=12, learning_rate=1000, n_iter=1000 and n_iter_early_exag=250.

Pegasus calculates a FIt-SNE, which are fast approximations of t-SNE embeddings, using the pyFIt-SNE package from Kluger lab [https://github.com/KlugerLab/pyFIt-SNE]. Pegasus uses the following FIt-SNE parameters by default: perplexity=30, early_exaggeration=12, learning_rate=1000, max_iter=1000, stop_early_exag_iter=250 and mom_switch_iter=250.

Pegasus calculates a UMAP based on the *k*-NN graph constructed by the HSNW algorithm, using the umap package from Leland Mclnnes [https://github.com/lmcinnes/umap]. Pegasus uses the following UMAP parameters by default: n_neighbors=15, min_dist=0.5, spread=1.0, n_epochs=250 and learning_rate=1.0.

Pegasus calculates an FLE using the forceatlas2 package, a Java package developed for scSVA^29^. The forceatlas2 package improved upon the Gephi’s ForceAtlas2^27^ Java code by providing more efficient parallelization^29^ and stops iterations if the average distance between FLE coordinates in adjacent iterations is no greater than *δ* or the maximum number of iterations *⊓g* is reached, where *δ* and *n_δ_* are user-specified parameters. Pegasus sets *δ* = 2.0 and *n_δ_* = 5000 by default.

#### Deep-learning-based visualization

To generate Net-* embeddings, we first subsample cells by local density. To estimate a proxy of the local density for each cell we denote as *d_i_* the distance from cell *i* to its *k*^th^ nearest neighbor in the calculated *k*-NN graph, where k is a user-provided parameter set to k = 25 by default. *d_i_* is inversely proportional to the local cell density such that *p_i_*, the probability of sampling cell *i*, is

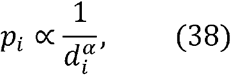

where *α* is a parameter that determines how much local density should influence the sampling process. If *α* = 0, we recover uniform sampling. In Pegasus, we set *α* = 1. We then subsample *P* percent of cells based on sampling probabilities {*p_i_*}, where *P* is a user-specified parameter set to 10% by default.

Next, we train a deep-learning-based regressor, using a neural network with 1 input layer and 4 hidden layers. The input layer connects with the top *m* principal components for each cell and the 4 hidden layers contain 100, 75, 50, and 25 ReLU units respectively. We train the network using scikit-learn’s MLPRegressor with a stochastic gradient decent solver, an adaptive learning rate and a *L*2 penalty parameter of 0.1. We additionally scale the inputs and outputs of the neural network, such that the maximal standard deviation of the inputs and outputs are 1.0 and 15.0 respectively.

In the final refinement step, Pegasus changes the following parameters for each embedding method. For t-SNE: learning rate 0.33 × *N*, where *N* is the total number of cells, n_iter=150 and n_iter_early_exag=0. For UMAP: learning_rate=10 and n_epochs=30. For FLE: *n_δ_* = 1500.

#### Differential expression analysis

Pegasus can perform Welch’s t-test, Fisher’s exact test and the Mann-Whitney U test between cells within and outside of a cluster. It controls the False Discovery Rate (FDR) at 5% using the Benjamini-Hochberg procedure^60^. Pegasus can optionally calculate the Area Under the ROC curve (AUROC) for each gene by considering the binary classification problem that uses the gene as the only feature to predict if a cell is within or outside of a cluster. Pegasus performs these analyses in parallel across genes to speed up the calculation process and outputs results for all genes in a spreadsheets. If AUROC is calculated, genes are ranked by their AUROC values.

#### Feature importance scores from LightGBM

Pegasus trains a LightGBM^30^ classifier on the log expression matrix to predict cluster labels. It uses 90% of the cells as the training data and 10% of the cells as the test data. It stops training the classifier once the prediction accuracy on the test dataset drops, and then extract each gene’s feature importance score from the trained classifier. To assign genes with high importance scores to each cluster, for each gene with a high importance score, it clusters the mean log expression levels of all clusters into 3 groups using the *k*-means algorithm, and assigns each of the 3 groups as strongly up-regulated, weakly up-regulated and down-regulated based on the mean log expression of the group. It then removes the group with largest size (as not significant) and assigns the gene to clusters in the other two groups.

#### Marker-based cell type annotation

To annotate cell types, Pegasus first loads known marker genes for each cell type from a user-provided JSON file. Each marker gene is associated with a sign (positive or negative) and a weight. The format of the JSON file is defined at [https://cumulus-doc.readthedocs.io/en/latest/cumulus.html#how-cell-type-annotation-works]. For each cluster, Pegasus enumerates all putative cell types and calculates a score between 0 and 1 per cell type, describing how likely cells from the cluster are to be of the specific cell type. To calculate the score, Pegasus assigns each marker a maximum impact value of 2. For a positive marker, if it is not up-regulated, its impact value is 0. Otherwise, if *fc* the fold change in the percentage of cells expressing this marker (within *vs*. outside of the cluster) satisfies *fc* ≥ 1.5, it has an impact value of 2 and is recorded as a strong supporting marker. If *fc* < 1.5, it has an impact value of 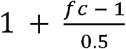 and is recorded as a weak supporting marker. For a negative marker, if it is up-regulated, its impact value is 0. If it is neither up-regulated nor down-regulated, its impact value 1 is 1. Otherwise, if 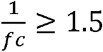, it has an impact value of 2 and is recorded as a strong supporting marker. If 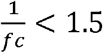, it has an impact value of 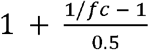 and is recorded as a weak supporting marker. The overall score is calculated as the ratio between the sum of impact values and the sum of weights multiplied by 2 from all expressed markers. Pegasus will evaluate all possible cell subtypes recursively if the current cell type score is no less than 0.5. Finally, Pegasus reports putative cell types with scores no less than *minimum_report_score* in descending order with respect to cell type scores for all clusters. By default, Pegasus sets *minimum_report_score* = 0.5.

### BENCHMARKING EXPERIMENTS

#### Benchmarked tools and benchmarking environments

We tested each component of Pegasus v0.15.0 and benchmarked it with SCANPY v1.4.4.post1 and Seurat v3.1.0 on key analysis tasks (**Supplementary Tables 1 and 2**) using a high-performance local server (Cloud-based times as part of a Cumulus workflow are in **Table 1**). The server has 28 CPU threads (1 Intel Xeon E5-2660v4 processor; 14-Core 2.00GHz, 35MB Cache) and 256 GB DDR4 ECC registered memory. In benchmarking, we used all 28 CPU threads whenever possible. For reproducibility, we have prepared a Docker image that has all three tools and their dependencies installed. This Docker image also contains instructions on how to reproduce the results shown in this manuscript. The Docker image is available at [https://hub.docker.com/r/cumulusprod/cumulus-experiment].

We also benchmarked Cumulus on cloud with running Cell Ranger + Seurat/SCANPY pipeline (on a 32 CPU-thread, 120GB Google Cloud virtual machine) on the bone marrow dataset. We used Cell Ranger v2.2.0 for this benchmark. The virtual machine for running Seurat (v3.1.0) and SCANPY (v1.4.4.post1) is created on Google compute engine zone us-west1-d, the same zone on which Cumulus ran its analysis step.

#### Bone marrow datasets pre-processing

We preprocessed the bone marrow data set by filtering out any cell with fewer than 500 genes or more than 6,000 genes, or where at least 10% of UMIs were from mitochondrial genes, retaining 274,182 cells. We then selected robust genes with *x* = 0.05%, normalized expression into TP100K and log-transformed the expression matrix.

#### HVG selection

We applied the standard and new HVG selection procedures to the log-transformed expression matrix separately, followed by the same downstream analyses on the resulting two sets of HVGs, using Pegasus with default parameters: batch correction, dimensionality reduction via PCA, *k*-NN graph construction, community detection using the Louvain algorithm, 2D visualization using FIt-SNE, differential expression analysis and marker-based cell type annotation.

We compared the highly variable genes selected using the two procedures with a list of immune genes curated by the ImmPort^34^ team (**Supplementary Fig. 1b**) from [https://www.immport.org/shared/geneData/GOappend1.xls], which contains 1,534 genes (with duplicates) annotated with immune-related gene ontology (GO) terms. The comparison results are available in **Supplementary Data 2**.

We evaluated the similarity between clusters obtained using the two HVG selection procedures with the adjusted mutual information^61^ (AMI) score defined below:

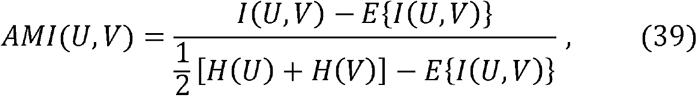

where *U* and *V* represent two cluster settings, *H* denotes entropy and *I* denotes mutual information.

#### Benchmarking of batch correction methods

We benchmarked Pegasus, ComBat, MNN, BBKNN and Seurat V3 using a subset of data from the bone marrow dataset, consisting of the first 10x Genomics channel from each of the 8 donors. Following the preprocessing steps above, we retained 34,654 cells. We then applied the new HVG selection procedure and downstream analyses described above (without batch correction) to obtain cell-type-annotated clusters (**Supplementary Fig. 2b**).

The clustering results showed one particular donor-specific effect (**Supplementary Fig. 2b**), with one donor-3-specific CD14^+^ monocyte cluster and one donor-3-specific T cell cluster. Since we do not want to count this donor-specific effect as “biology” when we compute kSIM acceptance rates, we constructed a ground truth for kSIM acceptance rates as follows. First, we merged the monocyte cluster into the larger monocyte cluster to its right. The donor-3-specific T cell cluster is adjacent to five other T cell clusters. We trained a LightGBM^30^ classifier that predicted cluster labels based on log expression levels using cells from the 5 clusters (90% training data + 10% validation data). The classifier’s accuracy on test data was 85.4%. We then used this classifier to assign each cell in the donor-3-specific T cell cluster into one of the 5 adjacent T cell clusters.

To generate batch-corrected results, we applied the same preprocessing step to the 34,654 cell dataset, selected top 2,000 HVGs using the new batch-aware HVG selection procedure, and extracted the HVG-specific gene-count matrix. With Pegasus, we applied the L/S adjustment method to the matrix to obtain batch-corrected expression levels. With SCANPY, we obtained ComBat-corrected and MNN-corrected expression levels. With Seurat v3 we obtained Seurat-corrected expression levels. We performed PCA, k-NN graph construction, and UMAP on the corrected expression matrices. With BBKNN, we used its output k-NN graph to replace Pegasus’s k-NN graph and kept other analyses the same.

We used kBET and kSIM acceptance rates on UMAP 2D coordinates. For kBET acceptance rate, we set *k* = 25 and *a* = 0.05. For kSIM acceptance rate, we set *k* = 25 and *β* = 0.9.

#### Benchmarking approximate nearest neighbor finding methods

We benchmarked the approximate nearest neighbor finding algorithms used by Pegasus, SCANPY and Seurat on the bone marrow dataset with default parameters. Pegasus ran the HNSW algorithm in full speed mode, SCANPY used the algorithm implemented in UMAP, and Seurat used the RcppAnnoy package at [https://cran.rstudio.com/web/packages/RcppAnnoy/index.html]. We ran the three methods on coordinates from the top 50 PCs produced by Pegasus and sought for top 100 nearest neighbors (including the cell itself). We also ran the brute force k-NN searching algorithm using scikit-learn^57^ to compute the ground truth. We evaluated each method’s performance using recall, defined as the percentage of *k* nearest neighbors that are also in the ground truth, and speed.

#### Diffusion pseudotime maps

We preprocessed the bone marrow dataset, selected HVGs using the new procedure, corrected batch effects, ran PCA and k-NN graph construction as described in HVG selection section. We generated diffusion pseudotime maps (with the parameters noted in **Fig. 2b** and **Supplementary Fig. 4**), visualized diffusion pseudotime maps using the FLE algorithm, and annotated the resulting trajectories using the cell type annotation described in the HVG selection section (with new HVG procedure).

#### Spectral community detection algorithms

We preprocessed the bone marrow dataset as described in HVG selection section, selected HVGs, corrected batch effects, ran PCA, k-NN graph construction, calculated a diffusion pseudotime map and performed 2D visualization using FIt-SNE with default parameters. We then generated different cluster settings using either the spectral clustering, Louvain, spectral-Louvain, Leiden, or spectral-Leiden algorithm. We performed differential expression analysis and marker-based cell type annotation for each of the clusterings separately.

#### Deep-learning-based visualization

We preprocessed the bone marrow dataset as described in HVG selection section, selected HVGs, corrected batch effects, ran PCA, k-NN graph construction, diffusion pseudotime map calculation using default parameters. We then ran Net-tSNE vs. t-SNE, Net-UMAP vs. UMAP and Net-FLE vs. FLE and annotated each visualization using cell types calculated in the HVG selection section (with the new HVG procedure).

#### Benchmarking Pegasus, SCANPY and Seurat on the full bone marrow dataset

We benchmarked Pegasus, SCANPY and Seurat on 10 tasks using the full bone marrow data of 274,182 cells. To ensure a fair comparison, whenever possible, all three methods received the same input computed using Pegasus with default parameters for each task. The only exception is Seurat, which does not accept a pre-computed affinity matrix as input for clustering. Thus, we provided Seurat with pre-computed principal components (PCs) instead and asked it to compute the affinity matrix before clustering. In addition, we used the *future* [https://cran.r-project.org/web/packages/future/] framework for parallelization as suggested at https://satijalab.org/seurat/v3.0/future_vignette.html. For the batch correction step, we made each 10x channel as a separate batch, which resulted in 63 batches in total. We used Seurat v3’s integration method^39^, BBKNN^38^ and L/S adjustment method^35^ to correct batch effects in Seurat, SCANPY and Pegasus, respectively. For *k*-NN graph construction, we set *k* = 100 for all three methods and set Pegasus in full speed mode. For Louvain-like and Leiden-like clustering, Pegasus used spectral-Louvain and spectral-Leiden algorithms, and Seurat and SCANPY used Louvain and Leiden algorithms. For t-SNE-like visualization, Seurat and Pegasus used FIt-SNE and SCANPY used Multicore-TSNE. For UMAP-like visualization, Pegasus used Net-UMAP, and Seurat and SCANPY used UMAP. We excluded the *k*-NN graph construction times for SCANPY and Pegasus, because it was accounted for in the *k*-NN graph construction task. Seurat calculates the *k*-NN graph again in the UMAP step using the *umap* Python package and thus we included *k*-NN graph construction time. For FLE-like visualization, Pegasus used Net-FLE and SCANPY used the fa2 package [https://github.com/bhargavchippada/forceatlas2]. The function calls, commands and parameters used for the three methods can be found in **Supplementary Note 2**.

#### Benchmarking Pegasus, SCANPY and Seurat v3 on the 1.3 million mouse brain dataset

We obtained the 1.3 million mouse brain data set from https://support.10xgenomics.com/single-cell-gene-expression/datasets/1.3.0/1M_neurons. The unfiltered data contains 1,306,127 cell barcodes. We preprocessed this data set by filtering out any cell with fewer than 500 genes or more than 6,000 genes, or with at least 10% of UMIs from mitochondrial genes. After filtration, we retained 1,286,072 cells. We selected robust genes with *x* = 0.05%, normalized expression into TP100K and log-transformed the expression matrix. Next, we benchmarked the three methods as described above. Since the fa2 package was too slow for our dataset, instead of running it for 5,000 iterations, we only ran it for 500 iterations and estimated total time by multiplying a factor of 10. The function calls, commands and parameters used for the three methods can be found in **Supplementary Note 2**.

#### Cloud computing execution time and cost

Cumulus utilizes Google Cloud Platform’s preemptible instances. Jobs running in preemptible instances can be kicked off by others’ jobs with higher priority but are 5x cheaper. By default, Cumulus allows up to 2 tries using preemptible instances before switching to non-kicked-off instances. Cumulus execution time is read out from Terra execution logs. To estimate the execution time of *mkfastq* and *count* steps on a 32 CPU-threads virtual machine (Table 1), we only sum over Terra-reported Docker running times of successful *mkfastq* and *count* runs, respectively. The *analysis* time for Seurat and SCANPY are estimated by running each tool on a same 32 CPU threads, 120 GB memory Google Cloud virtual machine instance. Since both Seurat and SCANPY do not have functions to aggregate 10x samples into a big count matrix, we used the aggregated matrix produced by Cumulus as their input and excluded the matrix aggregation time. In addition, since Seurat’s batch correction failed in the previous benchmarking (Fig. 2e), we reduced the number of batches from 63 to 8 (one per donor) and the number of PCs used from 30 (default) to 20. See Supplementary Note 2 for details on how Seurat, SCANPY and Cumulus analysis step were run. The total computational costs were reported by Terra and we calculated the average cost per sample by dividing 63.

