## Supplementary Data 2 for "Cumulus: a cloud-based data analysis framework for large-scale single-cell and single-nucleus RNA-seq"

**Cumulus Dependencies**

Cumulus code consists of four components: the Pegasus and scPlot python packages, the Cumulus WDL workflows and Dockfiles, the Cumulus docker images, and the Cirrocumulus app.

*The Pegasus python package depends on the following packages, each of which carries its own license and copyright:*

1.) anndata - BSD 3-Clause license (<https://github.com/theislab/anndata/blob/master/LICENSE>)

Copyright (c) 2017 - 2018 P. Angerer, F. Alexander Wolf, Theis Lab. All rights reserved.

2.) matplotlib - Matplotlib license (<https://matplotlib.org/users/license.html>)

Copyright (c) 2002 - 2012 John Hunter, Darren Dale, Eric Firing, Michael Droettboom and The Matplotlib Development Team; Copyright (c) 2012 - 2018 The Matplotlib Development team. All rights reserved.

3.) pandas - BSD 3-Clause license (<https://github.com/pandas-dev/pandas/blob/master/LICENSE>)

Copyright (c) 2008 - 2012 AQR Capital Management, LLC, Lambda Foundry, Inc. and PyData Development Team. All rights reserved.

4.) Cython - Apache 2.0 license (<https://github.com/cython/cython/blob/master/LICENSE.txt>)

Copyright (c) 2018 Cython contributors. All rights reserved.

5.) Scipy - BSD 3-Clause license (<https://scipy.org/scipylib/license.html>)

Copyright (c) 2001 - 2002 Enthought, Inc. All rights reserved.

Copyright (c) 2003 - 2013 SciPy Developers. All rights reserved.

6.) Seaborn - BSD 3-Clause license (<https://github.com/mwaskom/seaborn/blob/master/LICENSE>)

Copyright (c) 2012 - 2018 Michael L. Waskom. All rights reserved.

7.) scikit-learn - BSD 3-Clause license (<https://github.com/scikit-learn/scikit-learn/blob/master/COPYING>)

Copyright (c) 2007 – 2019 The scikit-learn developers. All rights reserved.

8.) statsmodels - BSD 3-Clause license (https://github.com/statsmodels/statsmodels/blob/master/LICENSE.txt)

Copyright (c) 2006 Jonathan E. Taylor. All rights reserved.

Copyright (c) 2006 - 2008 Scipy Developers. All rights reserved.

Copyright (c) 2009 - 2018 Statsmodels Developers. All rights reserved.

9.) natsort - MIT license (<https://github.com/bubkoo/natsort/blob/master/LICENSE>)

Copyright (c) 2016 W. Y. All rights reserved.

10.) numpy - BSD 3-Clause license (<https://docs.scipy.org/doc/numpy-1.10.4/license.html>)

Copyright (c) 2005 NumPy Developers. All rights reserved.

11.) tables – BSD 3-Clause license (<https://github.com/PyTables/PyTables/blob/master/LICENSE.txt>)

Copyright (c) 2002 - 2004 Francesc Alted. All rights reserved.

Copyright (c) 2005 - 2007 Carabos Coop. V. All rights reserved.

Copyright (c) 2008 - 2010 Francesc Alted. All rights reserved.

Copyright (c) 2011 - 2015 PyTables maintainers. All rights reserved.

12.) xlsxwriter - BSD license (<https://xlsxwriter.readthedocs.io/license.html>)

Copyright (c) 2013 John McNamara. All rights reserved.

13.) fisher - BSD license (<https://pypi.org/project/fisher/>)

Copyright (c) 2016 – 2019 Haibao Tang and Brent Pedersen. All rights reserved.

14.) loompy - BSD 2-Clause "simplified" license (<https://github.com/linnarsson-lab/loompy/blob/master/LICENSE>)

Copyright (c) 2016 Linnarsson Lab. All rights reserved.

15.) louvain - GPL v3 license (<https://github.com/vtraag/louvain-igraph/blob/master/LICENSE>)

Copyright (c) 2015 – 2017 Vincent Traag. All rights reserved.

16.) MulticoreTSNE - BSD 3-Clause license (<https://github.com/DmitryUlyanov/Multicore-TSNE/blob/master/LICENSE.txt>)

Copyright (c) 2016 – 2019 Dmitry Ulyanov. All rights reserved.

17.) docopt - MIT license (<https://github.com/docopt/docopt/blob/master/LICENSE-MIT>)

Copyright (c) 2012 Vladimir Keleshev. All rights reserved.

18.) setuptools - MIT license (<https://github.com/pypa/setuptools/blob/master/LICENSE>)

Copyright (c) 2016 Jason R. Coombs. All rights reserved.

19.) plotly - MIT license (<https://github.com/plotly/plotly.py/blob/master/LICENSE.txt>)

Copyright (c) 2016 – 2018 Plotly, Inc. All rights reserved.

20.) pybind11 - BSD license (<https://pypi.org/project/pybind11/>)

Copyright (c) 2016 Wenzel Jakob. All rights reserved.

21.) umap-learn - BSD 3-Clause license (<https://github.com/lmcinnes/umap/blob/master/LICENSE.txt>)

Copyright (c) 2017 Leland McInnes. All rights reserved.

22.) FIt-SNE - BSD, MIT licenses (<https://github.com/KlugerLab/FIt-SNE/blob/master/LICENSE.txt>)

Copyright (c) 2014 Laurens van der Maatan (Delft University of Technology). All rights reserved.

Copyright (c) 2019 George Linderman. All rights reserved.

23.) pyarrow - Apache 2.0 license (<https://github.com/apache/arrow/blob/master/LICENSE.txt>)

Copyright (c) 2016 – 2019 Arrow Contributors. All rights reserved.

24.) xgboost - Apache 2.0 license (<https://github.com/dmlc/xgboost/blob/master/LICENSE>)

Copyright (c) 2016 xgboost Contributors. All rights reserved.

25.) hnswlib - Apache 2.0 license (<https://github.com/nmslib/hnswlib/blob/master/LICENSE>)

Copyright (c) 2017 – 2019 Yury Malkov and other contributors. All rights reserved.

26.) Gephi - CDDL 1.0 license and GNU GPL v3 license (<https://github.com/gephi/gephi/blob/master/COPYING.txt>)

Copyright (c) 2012 Gephi Consortium. All rights reserved.

27.) leidenalg – GPL v3 license (<https://github.com/vtraag/leidenalg/blob/master/LICENSE>)

Copyright (c) 2017-2019 Vincent Traag. All rights reserved.

28.) forceatlas2-python – GPL v3 license (<https://github.com/klarman-cell-observatory/forceatlas2-python/blob/master/LICENSE>)

Copyright (c) 2019 Broad Institute. All rights reserved.

29.) scplot - BSD 3-Clause license (<https://github.com/klarman-cell-observatory/scPlot/blob/master/LICENSE>)

Copyright (c) 2019 Broad Institute. All rights reserved.

30.) scikit-misc – BSD 3-Clause license (<https://github.com/has2k1/scikit-misc/blob/master/LICENSE>)

Copyright (c) 2016 Hassan Kibirige. All rights reserved.

*The scPlot python package depends on the following packages, each of which carries its own license and copyright:*

1.) anndata - BSD 3-Clause license (<https://github.com/theislab/anndata/blob/master/LICENSE>)

Copyright (c) 2017 - 2018 P. Angerer, F. Alexander Wolf, Theis Lab. All rights reserved.

2.) numpy - BSD 3-Clause license (<https://docs.scipy.org/doc/numpy-1.10.4/license.html>)

Copyright (c) 2005 NumPy Developers. All rights reserved.

3.) Scipy - BSD 3-Clause license (<https://scipy.org/scipylib/license.html>)

Copyright (c) 2001 - 2002 Enthought, Inc. All rights reserved.

Copyright (c) 2003 - 2013 SciPy Developers. All rights reserved.

4.) pandas - BSD 3-Clause license (<https://github.com/pandas-dev/pandas/blob/master/LICENSE>)

Copyright (c) 2008 - 2012 AQR Capital Management, LLC, Lambda Foundry, Inc. and PyData Development Team. All rights reserved.

5.) hvplot - BSD 3-Clause license (<https://github.com/pyviz/hvplot/blob/master/LICENSE>)

Copyright (c) 2018, PyViz. All rights reserved.

6.) holoviews - BSD 3-Clause license (<https://github.com/pyviz/holoviews/blob/master/LICENSE.txt>)

Copyright (c) 2005 - 2019, IOAM (ioam.github.com). All rights reserved.

7.) colorcet - Creative Commons Attribution 4.0 International Public License (CC-BY) (<https://github.com/pyviz/colorcet/blob/master/LICENSE.txt>)

Copyright (c) 2019 colorcet contributors. All rights reserved.

*Cumulus dockers depend on the following software:*

1.) Cellranger - MIT license (<https://github.com/10XGenomics/cellranger/blob/master/LICENSE>)

Copyright (c) 2018 10X Genomics, Inc. All rights reserved.

2.) RSEM - GPL v3 license (<https://github.com/deweylab/RSEM/blob/master/COPYING>)

Copyright (c) 2011 Bo Li, Dewey Lab. All rights reserved.

3.) Bowtie 2 – GPL v3 license (<https://github.com/BenLangmead/bowtie2/blob/master/LICENSE>)

Copyright (c) 2011 – 2019 Ben Langmead and other Contributors. All rights reserved.

4.) Drop-seq Java tools – MIT license (<https://github.com/broadinstitute/Drop-seq/blob/master/LICENSE>)

Copyright (c) 2018 Broad Institute. All rights reserved.

5.) dropEst ­– GPL v3 license (<https://github.com/hms-dbmi/dropEst/blob/master/LICENSE.md>)

Copyright (c) 2018 Viktor Petukhov, Kharchenko Lab. All rights reserved.

*Cirrocumulus depends on the following software:*

1.) numpy - BSD 3-Clause license (<https://docs.scipy.org/doc/numpy-1.10.4/license.html>)

Copyright (c) 2005 NumPy Developers. All rights reserved.

2.) pandas - BSD 3-Clause license (<https://github.com/pandas-dev/pandas/blob/master/LICENSE>)

Copyright (c) 2008 - 2012 AQR Capital Management, LLC, Lambda Foundry, Inc. and PyData Development Team. All rights reserved.

3.) natsort - MIT license (<https://github.com/bubkoo/natsort/blob/master/LICENSE>)

Copyright (c) 2016 W. Y. All rights reserved.

4.) pyarrow - Apache 2.0 license (<https://github.com/apache/arrow/blob/master/LICENSE.txt>)

Copyright (c) 2016 – 2019 Arrow Contributors. All rights reserved.

5.) cachecontrol - Apache 2.0 license (<https://github.com/ionrock/cachecontrol/blob/master/LICENSE.txt>)

Copyright (c) 2015 Eric Larson. All rights reserved.

6.) gcsfs – BSD-style license (<https://github.com/dask/gcsfs/blob/master/LICENSE.txt>)

Copyright (c) 2017 Continuum Analytics, Inc. and contributors. All rights reserved.

7.) ujson - BSD-style license (<https://github.com/esnme/ultrajson/blob/master/LICENSE.txt>)

Copyright (c) 2014 Electronic Arts Inc. All rights reserved.

8.) flask - BSD 3-Clause license (<https://github.com/pallets/flask/blob/master/LICENSE.rst>)

Copyright (c) 2010 Pallets. All rights reserved.

9.) google-auth - Apache 2.0 license (<https://github.com/googleapis/google-auth-library-python/blob/master/LICENSE>)

Copyright (c) 2019 google-auth-library-python contributors. All rights reserved.

10.) google-cloud-datastore - Apache 2.0 license (<https://github.com/GoogleCloudPlatform/google-cloud-datastore/blob/master/LICENSE>)

Copyright (c) 2013 Google, Inc. and google-cloud-datastore contributors. All rights reserved.

*We also provide instructions for users to build their private bcl2fastq2 containing dockers. Bcl2fastq2 has the following license and endusers are responsible for compliance with that license:*

1.) Illumina's bcl2fastq2 - Illumina End User Software License (<https://support.illumina.com/content/dam/illumina-support/documents/downloads/software/bcl2fastq/bcl2fastq2-v2-20-eula.pdf>)

Copyright (c) 2017 Illumina, Inc. All rights reserved.
