## Supplemental Information for "Cumulus: a cloud-based data analysis framework for large-scale single-cell and single-nucleus RNA-seq"

Li B. et al.

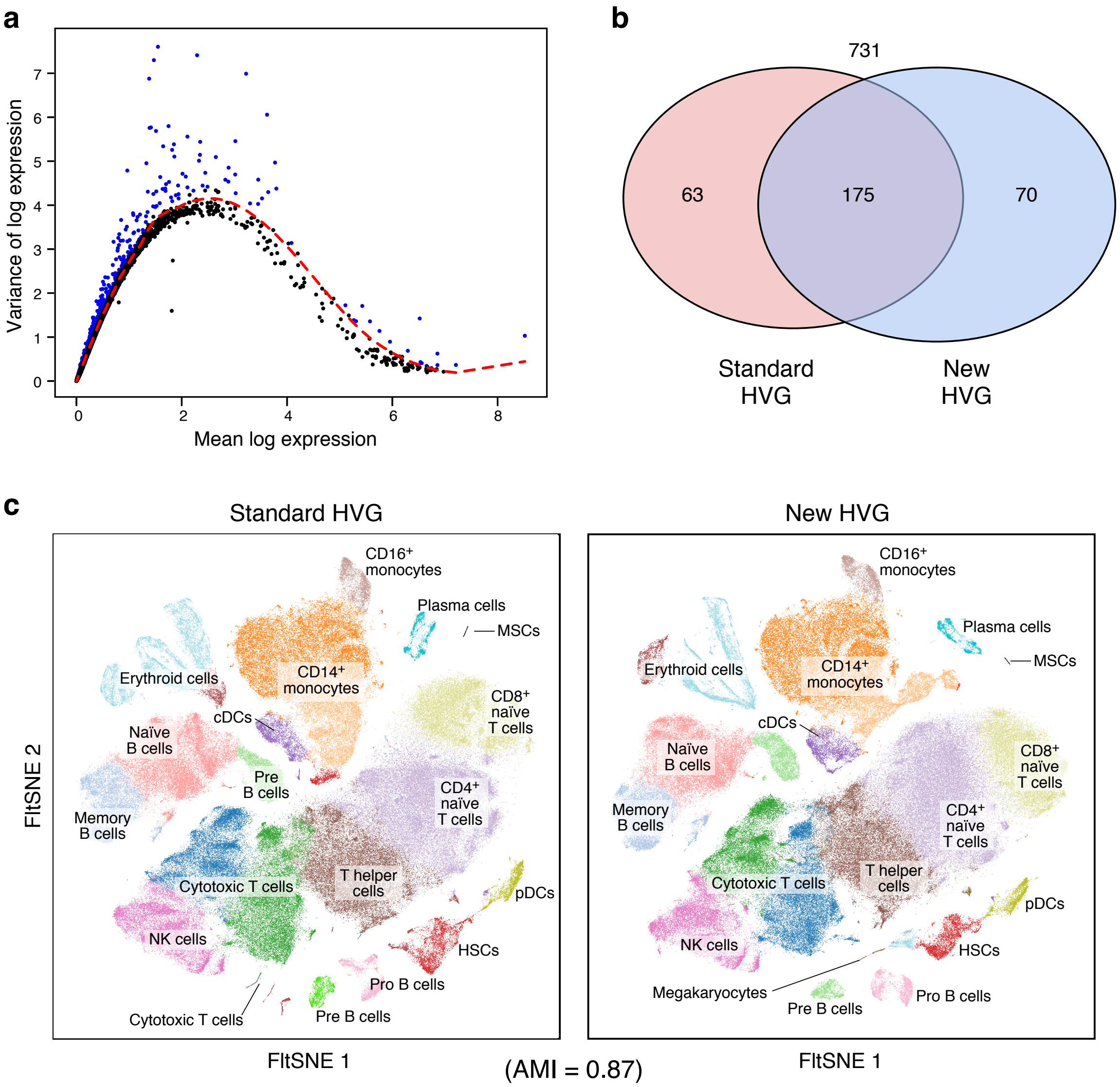


**Supplementary Figure 1. The new HVG selection procedure provides excellent quality *vs.* the standard procedure. a.** New HVG selection procedure. Variance (y axis) vs. mean (x axis) of log expression. Red: fit LOESS curve. HVGs (blue) are defined as the genes above the LOESS curve. **b.** Curated immune genes captured by each procedure. The number of ImmPort curated immune genes selected by a standard HVG procedure (red) and Cumulus (blue). **c.** Analysis with HVGs by new approach highlighted an additional cell type. FItSNE plots of cells from the bone marrow dataset generated by Cumulus with HVG genes selected by the standard (left) and new (right) procedure and colored by cell subset annotations. Bottom: Adjusted Mutual Information (AMI) score shows overall high concordance. Only the plot from the new procedure identifies megakaryocytes. HSCs: hematopoietic stem cells; MSCs: mesenchymal stem cells; cDCs: conventional dendritic cells; pDCs: plasmacytoid dendritic cells; NK cells: natural killer cells.

**a**

| | Pegasus | ComBat | MNN | BBKNN | Seurat v3 |
|---|---|---|---|---|---|
| **Batch correction time:** | 2 s | 20 s | 33 min, 55 s | 10 s | 8 min, 19 s |

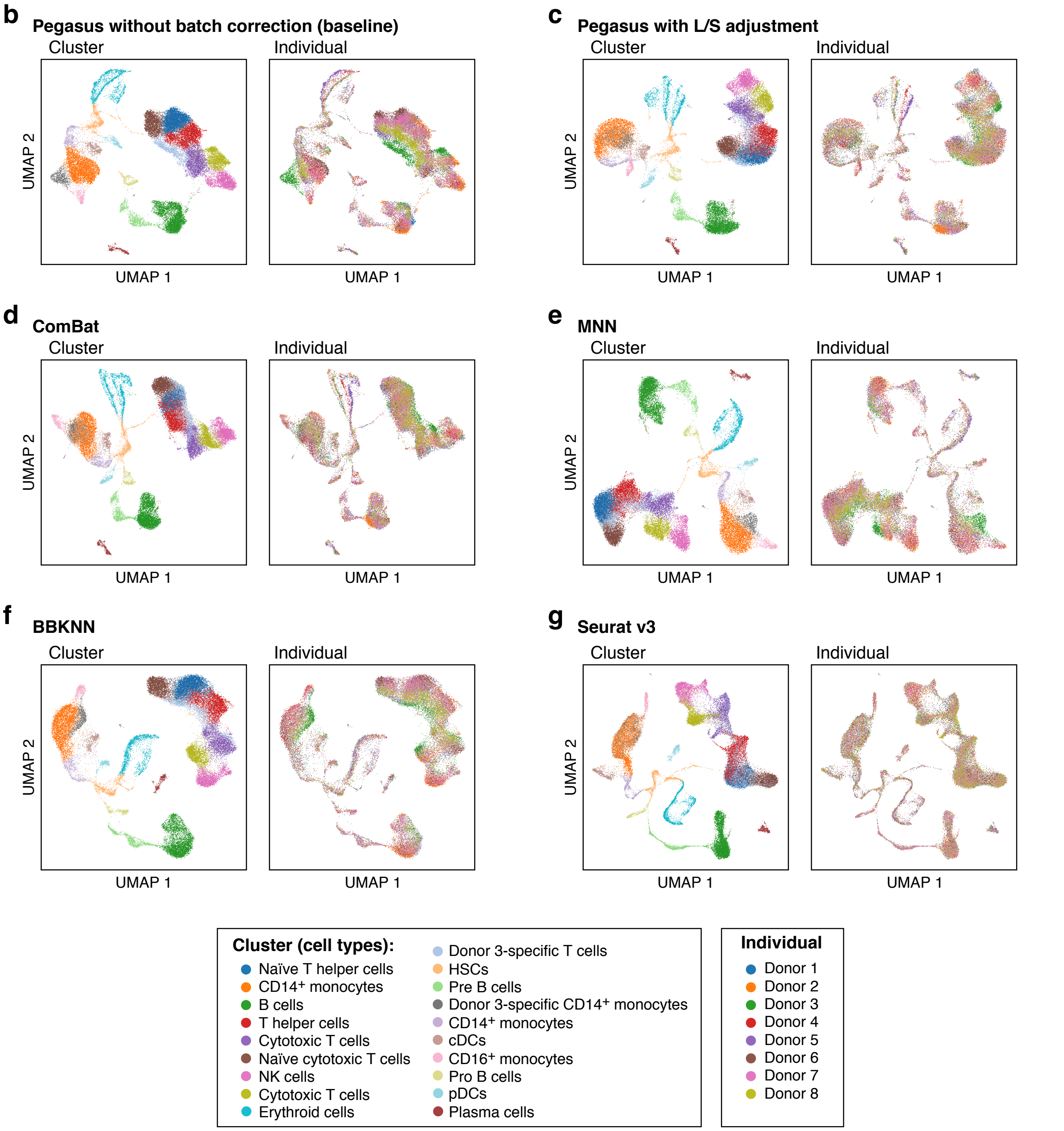


**Supplementary Figure 2. Benchmarking of batch correction methods on 34,654 bone marrow cells. a.** Execution time of each method. **a.-g.** UMAP visualizations of the bone marrow cells colored by either cell type annotation (left) or donor identity (right) without batch correction (**b**, baseline), Pegasus with L/S adjustment (**c**), ComBat (**d**), MNN (**e**), BBKNN (**f**), and Seurat v3 (**g**).

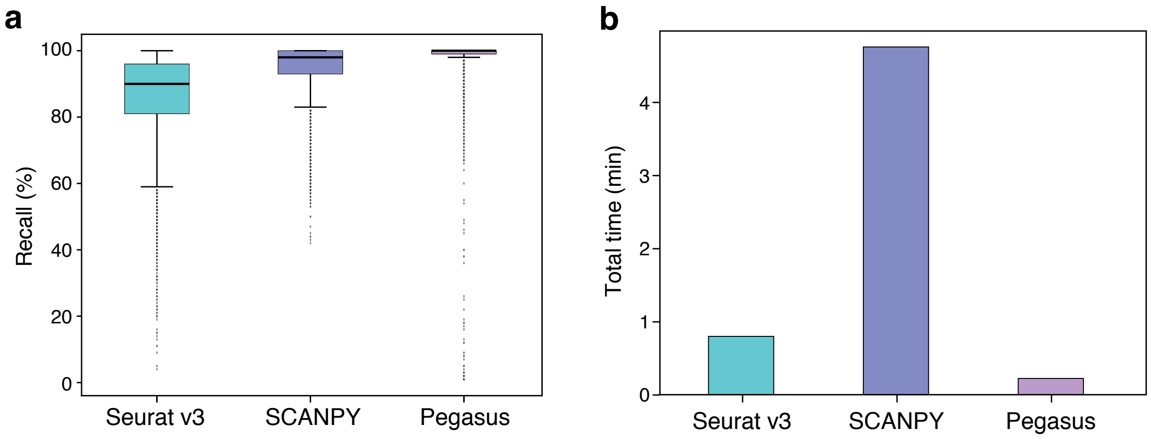


**Supplementary Figure 3. Benchmarking of approximate nearest neighbor finding methods on the bone marrow dataset.** Accuracy (**a**, y axis, % recall, **Methods**) and speed (**b**, y axis, minutes) of each of three methods. Boxplot (a): Line: median; box boundaries: lower and upper quartiles; whiskers: 1.5 interquartile range (IQR) below and above the low and high quartile, respectively.

a

b

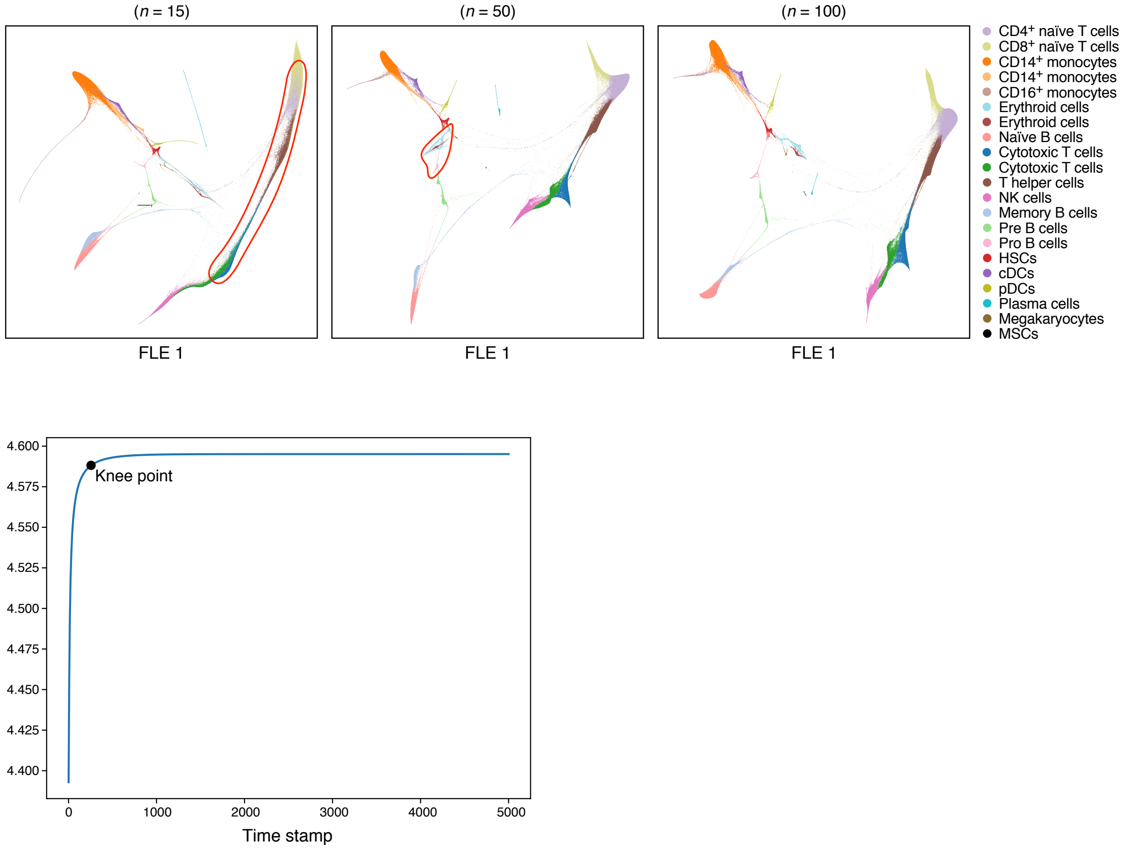


**Supplementary Figure 4. Adjusting diffusion pseudotime map parameters for visualization of pseudotemporal trajectories. a.** Using a large number of diffusion pseudotime components yields a developmental trajectory that enhances separation of trajectories of different cell populations. FLEs of single cell (colored by cell type annotation) generated from diffusion pseudotime maps ($t = \infty$) with 15 (left), 50 (middle) or 100 (right) components. $CD8^+$ and $CD4^+$ naïve T cells are fused together in the left FLE (circled in red). Erythrocytes and Pro-B cells are overlapped in the middle FLE (circled in Red). **b.** Choosing the timescale t for a diffusion pseudotime map. Von Neumann entropy (y axis) for diffusion maps with 100 components calculated from the bone marrow data at different timescales (x axis). Black point: knee point.

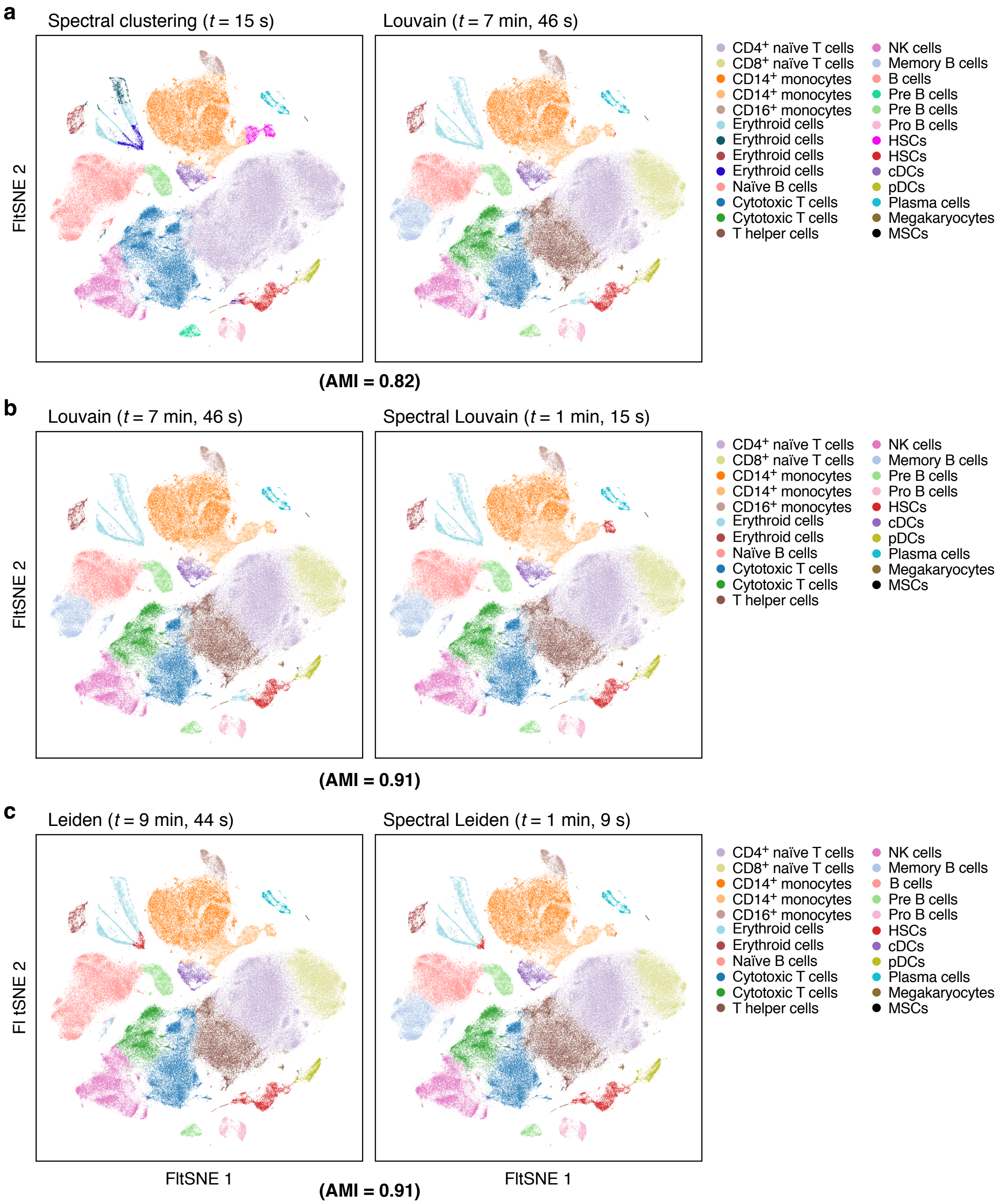


**Supplementary Figure 5. Spectral community detection algorithms combine the strengths of spectral clustering and community detection algorithms.** FIt-SNE of single cells (dots) colored by cluster assignment from (**a**) Spectral (left) *vs.* Louvain (right) clustering; (**b**) Louvain (left) *vs.* Spectral Louvain (right) clustering; or (**c**) Leiden (left) *vs.* spectral Leiden (right) clustering. Top: Execution time; bottom: Adjusted Mutual Information (AMI). *Post hoc* annotation labels are listed.

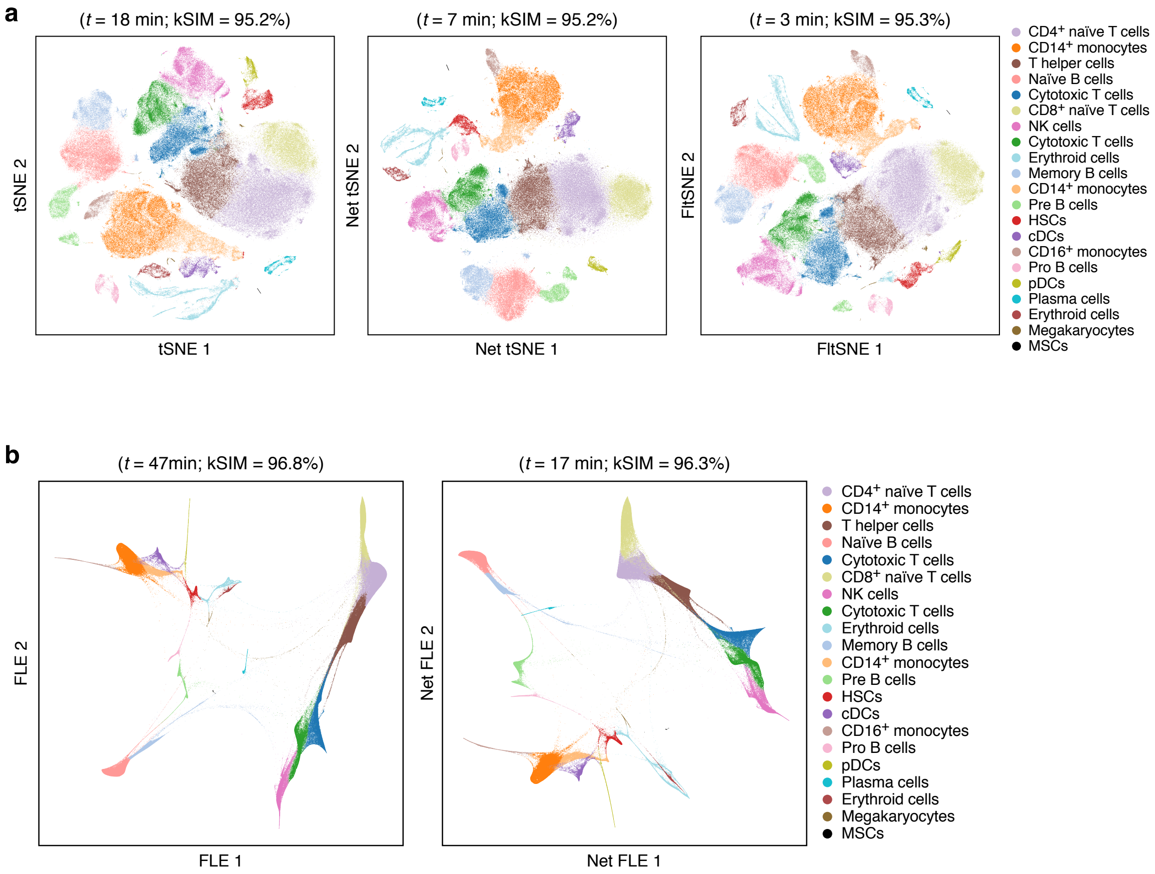


**Supplementary Figure 6. Deep-learning based visualization speeds up t-SNE and FLE visualizations while maintaining comparable quality.** Visualization of cell profiles (dots) from the full bone marrow data set colored by the same Louvain cluster membership (color; legend shows *post hoc* annotations) and laid out by (**a**) t-SNE (left), Net-tSNE (middle), or FIt-SNE (right); or (**b**) by FLE (left) or Net-FLE (right). Top: Execution time and kSIM acceptance rate.

| Feature | Cumulus | Cell Ranger | Seurat v3 | SCANPY |
|---|---|---|---|---|
| Cloud-based | ✓ | | | |
| Web-based user interface | ✓ | | | |
| Web-based interactive visualization | ✓ | | | |
| Reads extraction (from sequencers) | ✓ | ✓ | | |
| Count matrix generation: Droplet-based data (10x v2 and v3) | ✓ | ✓ | | |
| Count matrix generation: Droplet-based data (Drop-seq) | ✓ | | | |
| Count matrix generation: Plated-based data (SMART-Seq2) | ✓ | | | |
| Biological analyses | ✓ | ✓ | ✓ | ✓ |
| CITE-seq | ✓ | ✓ | ✓ | |
| Cell hashing | ✓ | | ✓ | |
| Nucleus hashing | ✓ | | | |
| Perturb-seq | ✓ | ✓ | | |

**Supplementary Table 1. Comparison of features (rows) between Cumulus and Cell Ranger, Seurat v3 and SCANPY (columns).**

| Feature | Pegasus | Seurat v3 | SCANPY |
|---|---|---|---|
| Quality control | ✓ | ✓ | ✓ |
| Highly variable gene selection | ✓ | ✓ | ✓ |
| Batch correction | ✓ | ✓ | ✓ |
| Data integration | | ✓ | ✓ |
| PCA | ✓ | ✓ | ✓ |
| $k$ nearest neighbor graph construction | ✓ | ✓ | ✓ |
| Diffusion map | ✓ | | ✓ |
| Pseudotime calculation | ✓ | | ✓ |
| Branching events detection | | | ✓ |
| Louvain-like clustering | ✓ | ✓ | ✓ |
| Leiden-like clustering | ✓ | ✓ | ✓ |
| tSNE-like visualization | ✓ | ✓ | ✓ |
| UMAP-like visualization | ✓ | ✓ | ✓ |
| FLE-like visualization | ✓ | | ✓ |
| Gene signature scores | | ✓ | ✓ |
| Differntial expression analysis | ✓ | ✓ | ✓ |
| Classification-based marker detection | ✓ | | |
| Marker-based cell type annotation | ✓ | | |
| Imputation | | | ✓ |
| Simulation | | | ✓ |
| Label transfer | | ✓ | |

**Supplementary Table 2. Comparison of features supported by Pegasus, Seurat v3 and SCANPY.**

| Analysis | Seurat v3 | SCANPY | Pegasus |
|---|---|---|---|
| HVG selection | 2 min, 7 s | 1 min, 27 s | **11 s** |
| Batch correction | Failed after 5h, 39 min | 8 min, 4 s | **19 s** |
| PCA | 5 min, 47 s | 12 s | **11 s** |
| Find $k$ nearest neighbors | 55 s | 4 min, 43 s | **15 sec** |
| Diffusion map | n/a | 4 min, 8 s | **2 min, 47 s** |
| Louvain-like clustering | 9 min, 32 s | 23 min, 48 s | **1 min, 15 s** |
| Leiden-like clustering | Failed | 43 min, 22 s | **1 min, 9s** |
| tSNE-like visualization | 3 min, 3 s | 19 min, 2 s | **2 min, 49 s** |
| UMAP-like visualization | 6 min, 28 s | 5 min, 49 s | **2 min, 24 s** |
| FLE-like visualization | n/a | 15 h, 36 min | **15 min, 20 s** |

**Supplementary Table 3. Execution time for 10 key analysis tasks on the bone marrow dataset by each tool.** Every tool was run on a 28-thread, 256G memory server. Bold font: shortest execution time.

| Analysis | SCANPY | Pegasus |
|---|---|---|
| HVG selection | 3 min, 43 s | **1 min, 50 s** |
| Batch correction | 1 h, 26 min | **2 min, 18 s** |
| PCA | **57 s** | **57 s** |
| Find $k$ nearest neighbors | 26 min, 10 s | **1 min, 28 s** |
| Diffusion map | 28 min, 4 s | **20 min, 13 s** |
| Louvain-like clustering | 1 h, 41 min | **7 min, 10 s** |
| Leiden-like clustering | 15 h, 59 min | **6 min, 24 s** |
| tSNE-like visualization | 1 h, 39 min | **11 min, 48 s** |
| UMAP-like visualization | 36 min, 31 s | **11 min, 28 s** |
| FLE-like visualization | $\approx$ 3 d 20 h | **1 h, 38 min** |

**Supplementary Table 4. Execution time for 10 key analysis tasks on the the 1.3 million mouse brain dataset by each tool.** Every tool was run on a 28-thread, 256G memory server. Seurat v3 failed to load the big gene-count matrix and thus is omitted from the table. Bold font: shortest execution time.

**Supplementary Data Captions**

**Supplementary Data 1. ImmPort-curated immune genes selected by the two HVG selection procedures.** Tab 1 (GOappend1_Immport): a copy of the original ImmPort spreadsheet from https://www.immport.org/shared/geneData/GOappend1.xls, Tab 2 (Duplicates_merged),: removed any duplicated genes; Tab 3 (Common_markers): ImmPort genes that were selected by both procedures, Tab 4 (Standard_procedure_specific) and Tab 5 (New_procedure_specific): ImmPort genes that are only selected by the standard procedure and the new procedure procedures, respectively.

**Supplementary Data 2. Cumulus dependencies and their licenses.**

**Supplementary Video Captions**

**Supplementary Video 1. Demonstration on how to submit Cumulus jobs on the Terra platform.**

**Supplementary Video 2. Demonstration of Cirrocumulus on the bone marrow dataset.**

### 1 Supplementary Note 1

#### 1.1 Preprocessing

The preprocessing step consists of 5 subtasks: *making gene symbols unique*, *selecting high quality cells*, *selecting robust genes*, *generating optional quality control reports*, and *transforming counts into log expression values*.

**Making gene symbols unique.** Pegasus takes a gene-by-cell count matrix as input. In this matrix, each gene is represented by a gene symbol. However, it is possible that multiple genes have a same gene symbol. To make sure each gene has a unique gene symbol, Pegasus appends suffix strings to genes with duplicated gene symbols. In particular, suppose that genes $1, \ldots, n$ have the same gene symbol "sym". Pegasus will transform gene $i$'s gene symbol from "sym" to "sym.$(i-1)$" for $i \geq 2$. (e.g. gene 2's symbole is "sym.1").

**Selecting high quality cells.** Pegasus enables users to select high quality cells based on a combination of the following criteria: (1) number of detected genes (with at least 1 Unique Molecular Identifier (UMI)) $\geq$ *min_genes*; (2) number of detected genes $<$ *max_genes*; (3) number of total UMIs $\geq$ *min_umis*; (4) number of total UMIs $<$ *max_umis*; (5) percentage of UMIs from mitochondrial genes $<$ *percent_mito*. *min_genes*, *max_genes*, *min_umis*, *max_umis*, and *percent_mito* are user-defined parameters.

**Selecting robust genes.** Genes that are only detected by a small portion of cells might correspond to random noise. Thus Pegasus identifies a set of robust genes which are detected in at least $x$ percent of total cells. Pegasus selects variable genes only from the robust gene set. $x$ is a user-defined parameter. The default value is $x = 0.05\%$, which corresponds to 3 cells out of $6,000$ cells.

**Generating optional quality control reports.** Pegasus can generate optional quality control reports as Excel spreadsheets and PDF figures.

The spreadsheet contains two tabs — `Cell filtration stats` and `Gene filtration stats`. `Cell filtration stats` tab contains 10 columns: `Channel`, `kept`, `median_n_genes`, `median_n_umis`, `median_percent_mito`, `filt`, `total`, `median_n_genes_before`, `median_n_umis_before`, and `percent_mito_before`. `Channel` lists names for channels included in the input data. `kept` lists the number of high quality cells each channel keeps. `median_n_genes`, `median_n_umis`, and `median_percent_mito` list the median number of detected genes, median number of UMIs and median percentage of mitochondrial UMIs among high quality cells for each channel. `filt` lists the number of low quality cells filtered out for each channel. `total` lists the total number of cells before filtration. `median_n_genes_before`, `median_n_umis_before`, and `percent_mito_before` list the median number of detected genes, median number of UMIs and median percentage of mitochondrial UMIs for each channel before filtration. `Gene filtration stats` tab has 3 columns: `gene`, `n_cells`, and `percent_cells`. `gene` lists gene symbols for all non-robust genes. `n_cells` lists the number of high quality cells each gene is detected. `percent_cells` lists the percentage of high quality cells having the gene detected. Genes are sorted in descending order with respect to `n_cells`.

Pegasus can also generates split violin plots in PDF format to compare the distribution of detected genes, total UMIs, or percentage of mitochondrial UMIs between high quality cells and all cells.

**Transforming counts into log expression values.** Because different cells are sequenced in different depth, we need to normalize each single cell so that their expression profiles are comparable [14]. Suppose we have $G$ genes in total and $R$ represents the set of robust genes. For a cell with UMI count vector $(c_1, \ldots, c_G)$, Pegasus multiplies a factor $\alpha$ to it to yield the normalized expression levels, $(y_1, \ldots, y_G)$:

$$(y_1, \ldots, y_G) = \alpha \cdot (c_1, \ldots, c_G), \quad \text{such that } \sum_{i \in R} \alpha c_i = c,$$

where $c$ is a user-defined constant. The default value is $c = 100,000$, which leads to expression levels in TP100K (transcripts per 100K).

Pegasus then transforms normalized expressions $(y_1, \ldots, y_G)$ into the log space. The log expression values $(Y_1, \ldots, Y_G)$ are calculated as follows:

$$Y_g = \log(y_g + 1). \tag{1}$$

#### 1.2 Highly variable gene selection

Highly variable gene (HVG) selection is a common feature selection step for scRNA-Seq data analysis. This step selects a set of informative genes showing higher than expected variances in their expression levels. Once we select HVGs, we perform all downstream analyses on these HVGs only.

Pegasus selects HVGs only among robust genes $R$ and implements two styles of selecting HVGs: Pegasus and Seurat.

##### 1.2.1 Pegasus style

We conduct all downstream analyses, such as dimension reduction and clustering, on the log expression space. It is natural to wonder why we do not select HVGs on the log expression space as well? Thus, Pegasus implements a simple HVG selection procedure directly on the log expression space.

**HVG selection in log expression space.** Suppose we have $N$ cells in total, we first calculate means and variances in the log space as follows for all genes in $R$:

$$\begin{aligned}
\widehat{\mu}_g &= \frac{1}{N}\sum_{i=1}^{N} Y_{ig}, \\
\widehat{\sigma}_g^2 &= \frac{1}{N-1}\sum_{i=1}^{N}(Y_{ig} - \widehat{\mu}_g)^2.
\end{aligned}$$

We then fit a LOESS curve [3] of degree 2 between the means and variances (Supplementary Fig. 1a) using scikit-misc [9] Python package. We set the span parameter to 0.02. We denote LOESS-predicted variance for gene $g$ as $\widetilde{\sigma}_g^2$ and we know that any gene $g$ with $\widehat{\sigma}_g^2 > \widetilde{\sigma}_g^2$ has a higher-than-expected variance.

There are two ways of ranking genes with higher-than-expected variances: by difference and by fold change. Let

$$\delta_g = \widehat{\sigma}_g^2 - \widetilde{\sigma}_g^2 \quad \text{and} \quad \tau_g = \frac{\widehat{\sigma}_g^2}{\widetilde{\sigma}_g^2}.$$

We rank each robust gene in $R$ in descending order with respect to $\delta_g$ and $\tau_g$ respectively and denote its rankings as $rank_\delta(g)$ and $rank_\tau(g)$. We then define the overall ranking of a gene as the sum of its two rankings to combine the strenghs of the two ranking methods:

$$rank(g) = rank_\delta(g) + rank_\tau(g).$$

HVGs are selected as the top $n$ genes according to $rank(g)$, where $n = 2000$ by default.

**Remove variance due to batch effects.** Batch effects are technical noises added to our data during the processes of sample handling, library preparation, and sequencing. We do not want batch effects to participate in the HVG selection procedure and thus need to come up a way to remove variances due to batch effects before we fit the LOESS curve.

Suppose we have $K$ biologically different groups, each group $k$ contains $n_k$ batches and each batch $j$ contains $n_{kj}$ cells. The total number of cells $N = \sum_{k=1}^{K}\sum_{j=1}^{n_k} n_{kj}$ and the mean and variance of each gene $g$ become

$$
\begin{aligned}
\widehat{\mu}_g &= \frac{1}{N}\sum_{k=1}^{K}\sum_{j=1}^{n_k}\sum_{i=1}^{n_{kj}} Y_{kjig}, \\
\widehat{\sigma}^2_g &= \frac{1}{N-1}\sum_{k=1}^{K}\sum_{j=1}^{n_k}\sum_{i=1}^{n_{kj}} (Y_{kjig} - \mu_g)^2.
\end{aligned}
$$

Based on the equations below, we can decompose the variance of gene $g$ ($\widehat{\sigma}^2_g$) into three components: variance within batches ($\widehat{\sigma}^2_{g1}$), variance between batches ($\widehat{\sigma}^2_{g2}$) and variance between biological groups ($\widehat{\sigma}^2_{g3}$).

$$
\begin{aligned}
\widehat{\sigma}^2_g &= \frac{1}{N-1}\sum_{k=1}^{K}\sum_{j=1}^{n_k}\sum_{i=1}^{n_{kj}} ((Y_{kjig} - \widehat{\mu}_{kjg}) + (\widehat{\mu}_{kjg} - \widehat{\mu}_{kg}) + (\widehat{\mu}_{kg} - \widehat{\mu}_g))^2, \\
&= \underbrace{\frac{1}{N-1}\sum_{k=1}^{K}\sum_{j=1}^{n_k}\sum_{i=1}^{n_{kj}} (Y_{kjig} - \widehat{\mu}_{kjg})^2}_{\widehat{\sigma}^2_{g1}} + \underbrace{\frac{1}{N-1}\sum_{k=1}^{K}\sum_{j=1}^{n_k} n_{kj}(\widehat{\mu}_{kjg} - \widehat{\mu}_{kg})^2}_{\widehat{\sigma}^2_{g2}} + \underbrace{\frac{1}{N-1}\sum_{k=1}^{K}(\sum_{j=1}^{n_k} n_{kj})\cdot(\widehat{\mu}_{kg} - \widehat{\mu}_g)^2}_{\widehat{\sigma}^2_{g3}},
\end{aligned}
$$

where

$$
\widehat{\mu}_{kjg} = \frac{1}{n_{kj}}\sum_{i=1}^{n_{kj}} Y_{kjig}, \quad \text{and} \quad \widehat{\mu}_{kg} = \frac{1}{\sum_{j=1}^{n_k} n_{kj}}\sum_{j=1}^{n_k}\sum_{i=1}^{n_{kj}} Y_{kjig}.
$$

We remove the variance due to batch effects by redefining $\widehat{\sigma}^2_g$ as

$$
\widehat{\sigma}^2_g := \widehat{\sigma}^2_{g1} + \widehat{\sigma}^2_{g3},
$$

and plug in the redefined variances to the previously described HVG selection procedure.

##### 1.2.2 Seurat style

Pegasus also implements the same HVG selection procedure that Seurat [14] and SCANPY [17] use.

**HVG selection in original expression space.** In this procedure, we first estimate mean, variance and index of dispersion of each gene $g$ in the original expression space ($y_{ig}$):

$$
\begin{aligned}
\widehat{\mu}_g &= \frac{1}{N}\sum_{i=1}^{N} y_{ig}, \\
\widehat{\sigma}^2_g &= \frac{1}{N-1}\sum_{i=1}^{N} (y_{ig} - \mu_g)^2, \\
\widehat{D}_g &= \frac{\widehat{\sigma}^2_g}{\widehat{\mu}_g}.
\end{aligned}
$$

We then transform the mean and index of dispersion into the log space:

$$
\begin{aligned}
\widehat{\mu}'_g &= \log(\widehat{\mu}_g + 1), \\
\widehat{D}'_g &= \log \widehat{D}_g.
\end{aligned}
$$

In the next step, We group genes into 20 bins based on their mean gene expression levels in the log space $\widehat{\mu}_g^{'}$ and calculates z-scores $z_g$ on log index of dispersion within each bin. Let us denote the set of genes in bin $j$ as $S_j$, we have

$$
\begin{aligned}
z_g &= \frac{\widehat{D}_g^{'} - \mu_j}{\sigma_j}, \\
\mu_j &= \frac{\sum_{g \in S_j} \widehat{D}_g^{'}}{|S_j|}, \\
\sigma_j &= \sqrt{\frac{1}{|S_j| - 1} \sum_{g \in S_j} (\widehat{D}_g^{'} - \mu_j)^2}.
\end{aligned}
$$

Now we can select HVGs based on the z-scores. Pegasus offers two ways to select HVGs: 1) Select top $n$ genes with largest z-scores, where $n = 2000$ by default; 2) Select all genes satisfying $a < \widehat{\mu}_g^{'} < b$ and $c < z_g < d$, where by default $a = 0.0125, b = 7, c = 0.5, d = \infty$. Note that in 2), all defaults are identical to SCANPY except $b$, where SCANPY's default is $b = 3$.

**Handle batch effects.** We adopt the method proposed in Seurat V3 [15] to handle batch effects during the HVG selection process. We first select top $n$ HVGs using the previous procedure independently for each batch. We then pool selected HVGs together and rank them by 1) number of batches the gene is in HVG set (descending order), and 2) the median rank in batches where the gene is HVG (ascending order). Lastly, we select the top $n$ genes from the pooled HVGs using the new ranking.

#### 1.3 Batch correction

Pegasus implements the simple location and scale (L/S) adjustment method [8, 10] for batch correction. Because the batch sizes in single-cell RNA-Seq data are very large in general (at least thousands of cells for droplet-based methods), there is no need to stabilize mean and variance estimates by using empirical Bayesian methods, such as the method used in ComBat [8]. We describe our correction method using mathmatical notations that are similar to [8].

**L/S method.** We first consider the scenario that we only have one biological group. Suppose we have $m$ batches and each batch $j$ has $n_j$ cells. We correct batch effects for all $G$ genes and we have $N$ cells, where $N = \sum_{j=1}^{m} n_j$. We model the log expression of gene $g$ in the $i$th cell of batch $j$ as

$$Y_{jig} = \alpha_g + \gamma_{jg} + \delta_{jg}\epsilon_{jig}, \qquad \epsilon_{jig} \sim Dist(0, \sigma_g^2),$$

where $\alpha_g$ is the average expression of gene $g$, and $\epsilon_{jig}$ is the error term following a distribution with 0 mean and $\sigma_g^2$ variance. $\gamma_{jg}$ and $\delta_{jg}$ are the additive and multiplicative batch effects for gene $g$ in batch $j$ respectively. In the L/S method, we adjust the batch effects for each gene separately.

**Parameter estimation.** Let $\ddot{\gamma}_{jg} = \alpha_g + \gamma_{jg}$, we have $Y_{jig} = \ddot{\gamma}_{jg} + \delta_{jg}\epsilon_{jig}$. The mean square error (MSE) estimate of $\ddot{\gamma}_{jg}$ is

$$\widehat{\ddot{\gamma}_{jg}} = \widehat{\alpha}_g + \widehat{\gamma}_{jg} = \frac{1}{n_j} \sum_{i=1}^{n_j} Y_{jig}.$$

To make sure the model is identifiable, we further impose the constraint that $\sum_j n_j \widehat{\gamma}_{jg} = 0$ [8]. With this constraint, we estimate $\alpha_g$ and $\gamma_{jg}$ as

$$
\begin{aligned}
\widehat{\alpha}_g &= \frac{1}{N} \sum_{j=1}^{m} \sum_{i=1}^{n_j} Y_{jig}, \\
\widehat{\gamma}_{jg} &= \frac{1}{n_j} \sum_{i=1}^{n_j} Y_{jig} - \widehat{\alpha}_g.
\end{aligned}
$$

Following [8], we estimate $\sigma_g$ as

$$\widehat{\sigma}_g = \sqrt{\frac{1}{N}\sum_{j=1}^{m}\sum_{i=1}^{n_j}(Y_{jig} - \widehat{\alpha}_g - \widehat{\gamma}_{jg})^2}.$$

Lastly, $\delta_{jg}$ can be estimated as

$$\widehat{\delta}_{jg} = \frac{\sqrt{\frac{1}{n_j-1}\sum_{i=1}^{n_j}(Y_{jig} - \widehat{\alpha}_g - \widehat{\gamma}_{jg})^2}}{\widehat{\sigma}_g}.$$

**Adjust batch effects.** Let us denote the adjusted expression leve as $Y^*_{jig}$, we have:

$$Y^*_{jig} = \frac{Y_{jig} - \widehat{\alpha}_g - \widehat{\gamma}_{jg}}{\widehat{\delta}_{jg}} + \widehat{\alpha}_g.$$

**Multiple biological groups.** Suppose we have $K$ groups of biological samples, each group $k$ has $n_k$ batches and each batch $kj$ has $n_{kj}$ cells. We further assume that any differences between the $K$ groups are biological. We model the log gene expression level as

$$Y_{kjig} = \alpha_{kg} + \gamma_{kjg} + \delta_{kjg}\epsilon_{kjig}, \qquad \epsilon_{kjig} \sim Dist(0, \sigma^2_{kg}),$$

adjust batch effects as

$$Y^*_{kjig} = \frac{Y_{kjig} - \widehat{\alpha}_{kg} - \widehat{\gamma}_{kjg}}{\widehat{\delta}_{kjg}} + \widehat{\alpha}_{kg}.$$

#### 1.4 Diffusion map and diffusion-based pseudotime

Pegasus implements the diffusion pseudotime (DPT) algorithm proposed in [6] with several improvements. To speed up the calculation of diffusion maps, we adopt 2 tricks that SCANPY [17] also uses: 1) constructing affinity matrix based on approximated kNN graph instead of the complete graph; 2) using only the top $n$ diffusion components to approximate diffusion distances, where $n$ is a user-specified parameter. In addition, based on diffusion maps, we define a family of diffusion pseudotime maps parameterized by timescale $t$, where the DPT algorithm corresponds to a diffusion pseudotime map with $t = \infty$. We provide a principled way of picking $t$ so that the corresponding diffusion pseudotime map reveals the embedded developmental trajectory better.

In the following, we first summarize how we compute diffusion maps. We then introduce how to calculate diffusion pseudotime maps and choose appropriate timescale $t$. Lastly, we describe how to approximate diffusion pseudotime maps with top $n$ components and how to calculate diffusion-based pseudotimes.

##### 1.4.1 Affinity matrix construction

Following [6], we put an isotropic Gaussian wave function on each cell $\mathbf{x}$, where $\mathbf{x} = (x_1, \ldots, x_n)^T$ represents the coordinates of this cell on top $n$ principal components:

$$Y_{\mathbf{x}}(\mathbf{x}') = \left(\frac{2}{\pi\sigma^2_{\mathbf{x}}}\right)^{\frac{1}{4}} \exp\left(-\frac{\|\mathbf{x}' - \mathbf{x}\|^2}{\sigma^2_{\mathbf{x}}}\right).$$

The kernel width $\sigma_{\mathbf{x}}$ is adjusted according to each cell $\mathbf{x}$. Suppose $d_1, \ldots, d_K$ are the distances between cell $\mathbf{x}$ and its $K$ nearest neighbors. Since $\mathbf{x}$'s 1st nearest neighbor is itself, $d_1 = 0$. We estimate $\sigma_{\mathbf{x}}$ as

$$\sigma_{\mathbf{x}} = \text{median}\,\{d_i \,|\, i = 2, \ldots, K\}.$$

The interference of the wave functions of two cells $\mathbf{x}$ and $\mathbf{y}$ results in a locally-scaled Gaussian kernel [5,18]:

$$
\begin{aligned}
K(\mathbf{x},\mathbf{y}) &= \int_{-\infty}^{\infty} Y_{\mathbf{x}}(\mathbf{x}')Y_{\mathbf{y}}(\mathbf{x}')d\mathbf{x}' \\
&= \int_{-\infty}^{\infty} \left(\frac{2}{\pi\sigma_{\mathbf{x}}^2}\right)^{\frac{1}{4}} \exp\left(-\frac{\|\mathbf{x}'-\mathbf{x}\|^2}{\sigma_{\mathbf{x}}^2}\right) \cdot \left(\frac{2}{\pi\sigma_{\mathbf{y}}^2}\right)^{\frac{1}{4}} \exp\left(-\frac{\|\mathbf{x}'-\mathbf{y}\|^2}{\sigma_{\mathbf{y}}^2}\right) d\mathbf{x}' \\
&= \int_{-\infty}^{\infty} \left(\frac{2}{\pi\sigma_{\mathbf{x}}\sigma_{\mathbf{y}}}\right)^{\frac{1}{2}} \exp\left(-\frac{\sigma_{\mathbf{y}}^2(\mathbf{x}'-\mathbf{x})^T(\mathbf{x}'-\mathbf{x})+\sigma_{\mathbf{x}}^2(\mathbf{x}'-\mathbf{y})^T(\mathbf{x}'-\mathbf{y})}{\sigma_{\mathbf{x}}^2\sigma_{\mathbf{y}}^2}\right) d\mathbf{x}' \\
&= \int_{-\infty}^{\infty} \left(\frac{2}{\pi\sigma_{\mathbf{x}}\sigma_{\mathbf{y}}}\right)^{\frac{1}{2}} \exp\left(-\frac{\sigma_{\mathbf{x}}^2+\sigma_{\mathbf{y}}^2}{\sigma_{\mathbf{x}}^2\sigma_{\mathbf{y}}^2}\left[\mathbf{x}'^T\mathbf{x}' - 2\left(\frac{\sigma_{\mathbf{y}}^2\mathbf{x}+\sigma_{\mathbf{x}}^2\mathbf{y}}{\sigma_{\mathbf{x}}^2+\sigma_{\mathbf{y}}^2}\right)^T\mathbf{x}' + \left(\frac{\sigma_{\mathbf{y}}^2\mathbf{x}+\sigma_{\mathbf{x}}^2\mathbf{y}}{\sigma_{\mathbf{x}}^2+\sigma_{\mathbf{y}}^2}\right)^T\left(\frac{\sigma_{\mathbf{y}}^2\mathbf{x}+\sigma_{\mathbf{x}}^2\mathbf{y}}{\sigma_{\mathbf{x}}^2+\sigma_{\mathbf{y}}^2}\right)\right]\right) \\
&\qquad\qquad \cdot \exp\left(-\frac{\mathbf{x}^T\mathbf{x}-2\mathbf{x}^T\mathbf{y}+\mathbf{y}^T\mathbf{y}}{\sigma_{\mathbf{x}}^2+\sigma_{\mathbf{y}}^2}\right) d\mathbf{x}' \\
&= \left(\frac{2\sigma_{\mathbf{x}}\sigma_{\mathbf{y}}}{\sigma_{\mathbf{x}}^2+\sigma_{\mathbf{y}}^2}\right)^{\frac{1}{2}} \exp\left(-\frac{\|\mathbf{x}-\mathbf{y}\|^2}{\sigma_{\mathbf{x}}^2+\sigma_{\mathbf{y}}^2}\right) \cdot \int_{-\infty}^{\infty} \left(\frac{1}{2\pi\frac{\sigma_{\mathbf{x}}^2\sigma_{\mathbf{y}}^2}{2(\sigma_{\mathbf{x}}^2+\sigma_{\mathbf{y}}^2)}}\right)^{\frac{1}{2}} \exp\left(-\frac{\|\mathbf{x}'-\frac{\sigma_{\mathbf{y}}^2\mathbf{x}+\sigma_{\mathbf{x}}^2\mathbf{y}}{\sigma_{\mathbf{x}}^2+\sigma_{\mathbf{y}}^2}\|^2}{2\frac{\sigma_{\mathbf{x}}^2\sigma_{\mathbf{y}}^2}{2(\sigma_{\mathbf{x}}^2+\sigma_{\mathbf{y}}^2)}}\right) d\mathbf{x}' \\
&= \left(\frac{2\sigma_{\mathbf{x}}\sigma_{\mathbf{y}}}{\sigma_{\mathbf{x}}^2+\sigma_{\mathbf{y}}^2}\right)^{\frac{1}{2}} \exp\left(-\frac{\|\mathbf{x}-\mathbf{y}\|^2}{\sigma_{\mathbf{x}}^2+\sigma_{\mathbf{y}}^2}\right).
\end{aligned}
$$

To construct the affinity matrix, we find the top $K$ nearest neighbors for each cell ($K = 100$ by default) using the state-of-the-art approximate nearest neighbor finding algorithm HNSW [11]. We then construct a sparse matrix $\mathbf{W}_1$ using the locally-scaled Gaussian kernel as follows:

$$
\mathbf{W}_1(i,j) = \begin{cases} K(\mathbf{x}_i,\mathbf{x}_j), & j \neq i \text{ and } j \text{ is in } i\text{'s } K \text{ nearest neighbors} \\ 0, & \text{otherwise} \end{cases}.
$$

Note that we exclude the 1st nearest neighbor of each cell, which is the cell itself, in the above equation.

We then obtain symmetrized matrix $\mathbf{W}_2$ by symmetrizing $\mathbf{W}_1$ as follows:

$$
\mathbf{W}_2(i,j) = \begin{cases} \mathbf{W}_1(i,j), & \mathbf{W}_1(i,j) > 0 \\ \mathbf{W}_1(j,i), & \mathbf{W}_1(i,j) = 0 \text{ and } \mathbf{W}_1(j,i) > 0 \\ 0, & \text{otherwise} \end{cases}.
$$

We assume the transcriptomic landscape is embedded in a low-dimensional manifold. Since there is no guarantee that the manifold is uniformly sampled and the sampling density is not necessarily related to the geometry of the manifold, we need to normalize each cell by its sampling density. Following [4], we first define the sampling density term $q(\mathbf{x}_i)$ for each cell $i$:

$$
q(\mathbf{x}_i) = \sum_{j=1}^{N} \mathbf{W}_2(i,j).
$$

We then define an anisotropic kernel $K'(\mathbf{x},\mathbf{y})$, which normalize the affinity between $\mathbf{x}$ and $\mathbf{y}$ by their sampling density terms:

$$
K'(\mathbf{x},\mathbf{y}) = \frac{K(\mathbf{x},\mathbf{y})}{q(\mathbf{x})q(\mathbf{y})}.
$$

We have an intuitive way to understand the above normalization. Suppose cell $i$ connects to cell $j$, which has a high sampling density and to cell $k$, which has a low sampling density. Then it is likely that $i$ connects to much more cells in $j$'s local neighborhood than $k$'s local neighborhood. After dividing the sampling density terms at $j$ and $k$, the total contributions of $j$'s neighborhood and $k$'s neighborhood to cell $i$ can be roughly equal.

By applying the anisotropic kernel to $\mathbf{W}_2$, we obtain the affinity matrix $\mathbf{W}$, which can recover the Riemannian geometry of the data, regardless of the sampling density [4]:

$$\mathbf{W}_{ij} = \frac{\mathbf{W}_2(i,j)}{q(\mathbf{x}_i)q(\mathbf{x}_j)}.$$

##### 1.4.2 Markov chain transition probability matrix

We define the Markov chain transition probability matrix $\mathbf{P}$ based on the affinity matrix $\mathbf{W}$:

$$\mathbf{P} = \mathbf{D}^{-1}\mathbf{W}, \quad \text{where } \mathbf{D} = diag(\sum_{j=1}^{N} \mathbf{W}_{ij}) \text{ is the degree matrix.}$$

The transition matrix $\mathbf{P}$ is not symmetric. We define the symmetric normalized transition matrix $\mathbf{Q}$ as

$$\mathbf{Q} = \mathbf{D}^{-\frac{1}{2}}\mathbf{W}\mathbf{D}^{-\frac{1}{2}},$$

and we have

$$\mathbf{P} = \mathbf{D}^{-\frac{1}{2}}\mathbf{Q}\mathbf{D}^{\frac{1}{2}}.$$

Since $\mathbf{Q}$ is real symmetric, it has the following eigen decomposition:

$$\mathbf{Q} = \mathbf{U}\mathbf{\Lambda}\mathbf{U}^T.$$

**Lemma 1.** *$\mathbf{Q}$'s eigenvalues $\lambda_i$ s are bounded between* $[-1,1]$.

*Proof.* We define $\mathbf{L}^{sym} = \mathbf{I} - \mathbf{Q}$, which is the symmetric normalized Laplacian matrix. $\mathbf{L}^{sym}$ is positive semidefinite because for any vector $\mathbf{c}$, we have

$$\frac{\mathbf{c}^T\mathbf{L}^{sym}\mathbf{c}}{\mathbf{c}^T\mathbf{c}} = \frac{\mathbf{c}^T(\mathbf{I}-\mathbf{Q})\mathbf{c}}{\mathbf{c}^T\mathbf{c}} = \frac{\sum_{i,j:i<j,\mathbf{Q}_{ij}>0} K'(\mathbf{x}_i,\mathbf{x}_j)\left(\frac{c_i}{\sqrt{d_i}} - \frac{c_j}{\sqrt{d_j}}\right)^2}{\|\mathbf{c}\|^2} \geq 0.$$

We denote $\mathbf{L}^{sym}$'s eigenvalues as $\lambda'_i$ s. We know $\mathbf{L}^{sym}$ and $\mathbf{Q}$ share the same eigenvectors and $\lambda_i = 1 - \lambda'_i$.

$$\begin{aligned} & \mathbf{L}^{sym}\mathbf{u}_i = \lambda'_i\mathbf{u}_i \\ \Leftrightarrow\ & (\mathbf{I}-\mathbf{Q})\mathbf{u}_i = \lambda'_i\mathbf{u}_i \\ \Leftrightarrow\ & \mathbf{Q}\mathbf{u}_i = (1-\lambda'_i)\mathbf{u}_i \\ \Leftrightarrow\ & \mathbf{Q}\mathbf{u}_i = \lambda_i\mathbf{u}_i. \end{aligned}$$

Since $\lambda'_i \geq 0$, we have $\lambda_i = 1 - \lambda'_i \leq 1$.

In addition, we can show $\lambda = 1$ and $\mathbf{u} = \mathbf{D}^{\frac{1}{2}}\mathbf{1}$ is an eigenvalue for $\mathbf{Q}$:

$$\mathbf{Q}\mathbf{u} = \mathbf{D}^{-\frac{1}{2}}\mathbf{W}\mathbf{D}^{-\frac{1}{2}} \cdot \mathbf{D}^{\frac{1}{2}}\mathbf{1} = \mathbf{D}^{-\frac{1}{2}} \cdot \mathbf{W}\mathbf{1} = \mathbf{D}^{-\frac{1}{2}}\mathbf{D}\cdot\mathbf{1} = \mathbf{D}^{\frac{1}{2}}\mathbf{1} = 1\mathbf{u}.$$

Similarly, we can show that $\mathbf{I}+\mathbf{Q}$ is also positive semidefinite since for any vector $\mathbf{c}$, we have

$$\frac{\mathbf{c}^T(\mathbf{I}+\mathbf{Q})\mathbf{c}}{\mathbf{c}^T\mathbf{c}} = \frac{\sum_{i,j:i<j,\mathbf{Q}_{ij}>0} K'(\mathbf{x}_i,\mathbf{x}_j)\left(\frac{c_i}{\sqrt{d_i}}+\frac{c_j}{\sqrt{d_j}}\right)^2}{\|\mathbf{c}\|^2} \geq 0.$$

Thus, $1+\lambda_i \geq 0 \Rightarrow \lambda_i \geq -1$.

Note that $\lambda_i = -1$ only if one of the connected component is a bipartite graph.

□

In practice, our single cell data always result in a connected graph. Thus, we assume eigenvalue $\lambda = 1$ only has one eigenvector. In addition, since we always observe cliques of at least 3 cells, it is unlikely the connected graph is a bipartite graph. Thus, we also assume that $\lambda_i > -1$.

We order $\mathbf{Q}$'s eigenvalues by their magnitudes: $1 = \lambda_0 > |\lambda_1| \geq |\lambda_2| \geq \cdots \geq |\lambda_{N-1}|$.

Now we can find $\mathbf{P}$'s eigen decomposition via $\mathbf{Q}$'s. We have

$$\begin{aligned}
\mathbf{P} &= \mathbf{D}^{-\frac{1}{2}}\mathbf{Q}\mathbf{D}^{\frac{1}{2}} \\
&= \mathbf{D}^{-\frac{1}{2}} \cdot \mathbf{U}\mathbf{\Lambda}\mathbf{U}^T \cdot \mathbf{D}^{\frac{1}{2}} \\
&= (\mathbf{D}^{-\frac{1}{2}}\mathbf{U})\mathbf{\Lambda}(\mathbf{D}^{\frac{1}{2}}\mathbf{U})^T \\
&= \mathbf{\Psi}\mathbf{\Lambda}\mathbf{\Phi}^T,
\end{aligned}$$

where $\mathbf{\Psi} = \mathbf{D}^{-\frac{1}{2}}\mathbf{U}$ are $\mathbf{P}$'s right eigenvectors and $\mathbf{\Phi} = \mathbf{D}^{\frac{1}{2}}\mathbf{U}$ are $\mathbf{P}$'s left eigenvectors. In addition, we have

$$\mathbf{\Phi}^T\mathbf{\Psi} = \mathbf{U}^T\mathbf{D}^{\frac{1}{2}} \cdot \mathbf{D}^{-\frac{1}{2}}\mathbf{U} = \mathbf{U}^T\mathbf{U} = \mathbf{I}.$$

Note that the right eigenvector of $\lambda_0 = 1$ for $\mathbf{P}$ becomes

$$\psi_0 = \mathbf{D}^{-\frac{1}{2}}\mathbf{u}_0 = \mathbf{D}^{-\frac{1}{2}} \cdot \mathbf{D}^{\frac{1}{2}}\mathbf{1} = \mathbf{1}.$$

##### 1.4.3 Diffusion maps and diffusion distances

We define a family of diffusion maps $\{\mathbf{\Psi}_t\}_{t\in\mathbb{N}}$ as

$$\mathbf{\Psi}_t(\mathbf{x}_i) = \begin{pmatrix} \lambda_1^t\psi_1(i) \\ \lambda_2^t\psi_2(i) \\ \vdots \\ \lambda_{N-1}^t\psi_{N-1}(i) \end{pmatrix},$$

and associated diffusion distances as

$$D_t(\mathbf{x}_i,\mathbf{x}_j) = \|\mathbf{\Psi}_t(\mathbf{x}_i) - \mathbf{\Psi}_t(\mathbf{x}_j)\| = \left(\sum_{l=1}^{N-1} \lambda_l^{2t}(\psi_l(i)-\psi_l(j))^2\right)^{\frac{1}{2}}.$$

In above definitions, we omit diffusion component 0, $\Psi_0$, since all its components are the same and thus have no influence on the distance.

We can show that the diffusion distance $D_t$ is equal to the weighted $\ell^2$ distance between transition probabilities after $t$ steps.

Let us define $p_t(\cdot,\cdot)$ as the transition probabilities between any two cells after $t$ steps:

$$
\begin{aligned}
p_t(\cdot,\cdot) &= \mathbf{P}^t \\
&= \mathbf{\Psi}\mathbf{\Lambda}\mathbf{\Phi}^T \cdot \mathbf{\Psi}\mathbf{\Lambda}\mathbf{\Phi}^T \cdots \mathbf{\Psi}\mathbf{\Lambda}\mathbf{\Phi}^T \\
&= \mathbf{\Psi}\mathbf{\Lambda}^t\mathbf{\Phi} \\
&= \sum_{l=0}^{N-1} \lambda_l^t \psi_l \phi_l^T.
\end{aligned}
$$

**Lemma 2.**

$$
D_t(\mathbf{x}_i, \mathbf{x}_j) = \|p_t(\mathbf{x}_i, \cdot) - p_t(\mathbf{x}_j, \cdot)\|_{\ell^2(\mathbb{R}^N, \frac{1}{d})}
$$

*Proof.*

$$
\begin{aligned}
\|p_t(\mathbf{x}_i,\cdot) - p_t(\mathbf{x}_j,\cdot)\|^2_{\ell^2(\mathbb{R}^N,\frac{1}{d})} &= \sum_{k=1}^{N} (\mathbf{P}_{ik}^t - \mathbf{P}_{jk}^t)^2 \cdot \frac{1}{d_k} \\
&= \sum_{k=1}^{N} \left( \sum_{l=0}^{N-1} \lambda_l^t \psi_l(i)\phi_l(k) - \sum_{l=0}^{N-1} \lambda_l^t \psi_l(j)\phi_l(k) \right)^2 \cdot \frac{1}{d_k} \\
&= \sum_{k=1}^{N} \left( \sum_{l=0}^{N-1} \lambda_l^t (\psi_l(i) - \psi_l(j))\phi_l(k) \right)^2 \cdot \frac{1}{d_k} \\
&= \sum_{k=1}^{N} \sum_{l=0}^{N-1} \sum_{l'=0}^{N-1} \lambda_l^t \lambda_{l'}^t (\psi_l(i) - \psi_l(j))(\psi_{l'}(i) - \psi_{l'}(j)) \frac{\phi_l(k)\phi_{l'}(k)}{d_k} \\
&= \sum_{l=0}^{N-1} \sum_{l'=0}^{N-1} \lambda_l^t \lambda_{l'}^t (\psi_l(i) - \psi_l(j))(\psi_{l'}(i) - \psi_{l'}(j)) \frac{\sum_{k=1}^{N} \phi_l(k)\phi_{l'}(k)}{d_k} \\
&= \sum_{l=0}^{N-1} \lambda_l^{2t} (\psi_l(i) - \psi_l(j))^2 \\
&= \sum_{l=1}^{N-1} \lambda_l^{2t} (\psi_l(i) - \psi_l(j))^2 \\
&= D_t(\mathbf{x}_i, \mathbf{x}_j)^2.
\end{aligned}
$$

□

A diffusion map $\mathbf{\Psi}_t$ with small $t$ reveals local geometric properties and is less robust to noise. In contrast, $\mathbf{\Psi}_t$ with large $t$ reveals global geometric properties and is more robust.

##### 1.4.4 Diffusion pseudotime maps

We define a family of diffusion pseudotime maps $\{\mathbf{\Psi}_t^{'}\}_{t\in\mathbb{N}}$ as

$$
\begin{aligned}
\mathbf{\Psi}_t^{'}(\mathbf{x}_i) &= \sum_{t'=1}^{t} \mathbf{\Psi}_{t'}(\mathbf{x}_i) \\
&= \begin{pmatrix} \sum_{t'=1}^{t} \lambda_1^{t'} \psi_1(i) \\ \sum_{t'=1}^{t} \lambda_2^{t'} \psi_2(i) \\ \vdots \\ \sum_{t'=1}^{t} \lambda_{N-1}^{t'} \psi_{N-1}(i) \end{pmatrix} \\
&= \begin{pmatrix} \lambda_1 \frac{1-\lambda_1^t}{1-\lambda_1} \psi_1(i) \\ \lambda_2 \frac{1-\lambda_2^t}{1-\lambda_2} \psi_2(i) \\ \vdots \\ \lambda_{N-1} \frac{1-\lambda_{N-1}^t}{1-\lambda_{N-1}} \psi_{N-1}(i) \end{pmatrix}.
\end{aligned}
$$

The associated diffusion pseudotime distances are

$$D_t^{'}(\mathbf{x}_i, \mathbf{x}_j) = \|\mathbf{\Psi}_t^{'}(\mathbf{x}_i) - \mathbf{\Psi}_t^{'}(\mathbf{x}_j)\| = \left(\sum_{l=1}^{N-1} [\lambda_l \frac{1-\lambda_l^t}{1-\lambda_l} \cdot (\psi_l(i) - \psi_l(j))]^2\right)^{\frac{1}{2}}.$$

If we set $t = \infty$, we recover the DPT algorithm [6], which sums diffusion maps over all timescales. However, when the timescale $t$ becomes large enough, the correpending diffusion map begins to smooth out real signals [12, 16]. In the extreme case $t = \infty$, all signals are smoothed out and the diffusion distance between any pair of cells is 0. Thus, $t = \infty$ might lose important biological signals. In Pegasus, we seek to pick a $t$ that smooth out most noise but little signal.

**von Neumann Entropy.** Inspired by the PHATE [12] work, we choose $t$ based on the von Neumann entropy [1] of the graph induced by each timescale. For each timescale $t$, the power matrix $\mathbf{P}^t$ induces a graph. Since $\mathbf{P}^t$ is a transition matrix, its degree matrix $\mathbf{D}$ is always equal to the identity matrix $\mathbf{I}$, and its Laplacian matrix becomes

$$\begin{aligned}
\mathbf{L}(t) &= \mathbf{I} - \mathbf{P}^t \\
&= \mathbf{\Psi}\mathbf{\Phi}^T - \mathbf{\Psi}\mathbf{\Lambda}^t\mathbf{\Phi}^T \\
&= \mathbf{\Psi}(\mathbf{I} - \mathbf{\Lambda}^t)\mathbf{\Phi}^T.
\end{aligned}$$

Thus $\mathbf{L}(t)$ shares the same eigenvectors as $\mathbf{P}$ and have eigenvalues $1 - \lambda_0^t, \ldots, 1 - \lambda_{N-1}^t$.

We then derive $\mathbf{P}^t$'s graph density matrix $\rho(\mathbf{L}(t))$ by normalizing $\mathbf{L}(t)$'s trace to 1:

$$\rho(\mathbf{L}(t)) = \frac{\mathbf{L}(t)}{tr(\mathbf{L}(t))}, \qquad \text{where } tr(\mathbf{L}(t)) = \sum_{i=0}^{N-1} 1 - \lambda_i^t.$$

Let us denote the density matrix's eigenvalues as $[\eta(t)]_0, \ldots, [\eta(t)]_{N-1}$, we have

$$[\eta(t)]_i = \frac{1 - \lambda_i^t}{\sum_{j=0}^{N-1} 1 - \lambda_j^t} \qquad \text{and} \qquad \sum_{i=0}^{N-1} [\eta(t)]_i = 1.$$

The von Neumann entropy $S(t)$ is defined on the density matrix $\rho(\mathbf{L}_t)$:

$$\begin{aligned}
S(t) &= -tr(\rho(\mathbf{L}(t)) \log \rho(\mathbf{L}(t))) \\
&= -\sum_{i=0}^{N-1} [\eta(t)]_i \log[\eta(t)]_i.
\end{aligned}$$

$S(t)$ increases as $t$ increases and reaches its maximum $S(t) = \log(N-1)$ when $t \to \infty$. As $t$ increases, the smaller eigenvalues (likely representing noise) of transition matrix $\mathbf{P}$ decreases to 0 (and eigenvalue of 1 for $\mathbf{L}(t)$) much more rapidly than the larger eigenvalues (likely representing signals) [12]. Thus, we can observe a high rate of increase in $S(t)$ initially when data noise is reduced and then a low rate of increase in $S(t)$ when signal is smoothed out.

If we plot $S(t)$ against $t$, we can see a knee-shaped curve and we pick the best $t$ as the knee point of this curve.

**Picking knee point.** To find the knee point, we consider all $t$s in the range of $[1, maxt]$, where $maxt$ is a user-specified parameter (default: $maxt = 5000$). We draw a segment between $t = 1$ and $t = maxt$ and find the point in curve that is furthest from the segment by scanning $t = 2, \dots, maxt - 1$.

We define $p_1 = \begin{bmatrix} 1 \\ S(1) \end{bmatrix}$ and $p_1 = \begin{bmatrix} maxt \\ S(maxt) \end{bmatrix}$ as the two end points, and let $p_3 = \begin{bmatrix} t \\ S(t) \end{bmatrix}$ as the point we scan. We can calculate the distance between $p_3$ and segment $p_2 - p_1$ as

$$dis = \frac{\|(p_3 - p_1) \times (p_2 - p_1)\|}{\|p_2 - p_1\|}.$$

##### 1.4.5 Approximation

Diffusion components with eigenvalues close to 0 contribute little to the diffusion distance. This implies that we can approximate the distance by only keeping the top $n$ diffusion components. Since we exclude diffusion component 0 ($\Psi_0$), we keep $n - 1$ diffusion components in practice. Pegasus uses $n = 100$ by default.

**Compute top $n$ eigenvalues and eigenvectors.** We use the Implicitly Restarted Lanczos Method [2] (via scipy.sparse.linalg.eigsh function) to calculate the top $n$ eigenvalues and eigenvectors of $\mathbf{Q}$.

We also provide an option to compute top $n$ eigenvalues and eigenvectors using the randomized SVD algorithm [7] implemented in scikit-learn [13] as follows:

Suppose $\mathbf{Q}$'s SVD decomposition is as follows:

$$\mathbf{Q} \approx \mathbf{U}_{N\times n} \cdot \mathbf{S}_{n\times n} \cdot \mathbf{V}^T_{n\times N}.$$

We know that $\mathbf{U}_{N\times n}$ are the top $n$ eigenvectors of $\mathbf{Q}$. However, the singular values are not necessarily equal to the eigenvalues. In particular, we have

$$\sigma_i = \sqrt{\lambda_i^2} = |\lambda_i|,$$

which means the singular values lose the signs.

We use $\mathbf{V}^T_{n\times N}$ to help us recover the signs. We know that

$$\mathbf{v}_i = \frac{\mathbf{Q}^T \mathbf{u}_i}{\sigma_i} = \frac{\lambda_i \mathbf{u}_i}{\sigma_i} = \text{sign}(\lambda_i)\mathbf{u}_i,$$

and thus

$$\mathbf{V}^T_{n\times N} \cdot \mathbf{U}_{N\times n} = \begin{pmatrix} \text{sign}(\lambda_0) & 0 & \cdots & 0 \\ 0 & \text{sign}(\lambda_1) & \cdots & 0 \\ \vdots & \vdots & \ddots & \vdots \\ 0 & 0 & \cdots & \text{sign}(\lambda_{n-1}) \end{pmatrix}.$$

Therefore, we can calculate eigenvalues $\mathbf{\Lambda}_{n\times n}$ as

$$\mathbf{\Lambda}_{n\times n} = (\mathbf{V}^T_{n\times N} \mathbf{U}_{N\times n}) \cdot \mathbf{S}_{n\times n}.$$

**von Neumann entropy for top $n$ eigenvalues.** If we only focus on the top $n$ eigenvalues, we obtain the following truncated Laplacian matrix:

$$\widetilde{\mathbf{L}}(t) = \sum_{i=0}^{n-1} (1 - \lambda_i^t) \psi_i \phi_i^T,$$

and its density matrix:

$$\rho(\widetilde{\mathbf{L}}(t)) = \frac{\widetilde{\mathbf{L}}(t)}{tr(\widetilde{\mathbf{L}}(t))}, \quad \text{where } tr(\widetilde{\mathbf{L}}(t)) = \sum_{i=0}^{n-1} 1 - \lambda_i^t.$$

Thus, the von Neumann entropy becomes

$$S(t) := -\sum_{i=0}^{n-1} [\eta'(t)]_i \log[\eta'(t)]_i, \quad \text{where } [\eta'(t)]_i = \frac{1 - \lambda_i^t}{\sum_{j=0}^{n-1} 1 - \lambda_j^t}.$$

###### 1.4.6 Diffusion-based pseudotime calculation

Suppose users select a set of $m$ cells as roots, we first calculate the diffusion-based distance for each cell as follows:

$$dis(\mathbf{x}_i, roots) = \frac{1}{m} \sum_{j \in roots} D_t^{'}(\mathbf{x}_i, \mathbf{x}_j) \approx \frac{1}{m} \sum_{j \in roots} \left( \sum_{l=1}^{n-1} [\lambda_l \frac{1 - \lambda_l^t}{1 - \lambda_l} \cdot (\psi_l(i) - \psi_l(j))]^2 \right)^{\frac{1}{2}}.$$

Then we normalize these distances to between $[0, 1]$ to obtain our diffusion-based pseudotimes:

$$pseudotime(\mathbf{x}_i) = \frac{dis(\mathbf{x}_i, roots) - \min_j dis(\mathbf{x}_j, roots)}{\max_j dis(\mathbf{x}_j, roots) - \min_j dis(\mathbf{x}_j, roots)}.$$

### 2 Supplementary Note 2

#### 2.1 Function calls, commands and parameters used for benchmarking Pegasus, SCANPY and Seurat on the full bone marrow dataset (Table S2)

**Seurat.** We first set `plan("multiprocess", workers=28)`. Next, we ran `SelectIntegrationFeatures` with parameters `mean.cutoff=c(0.0125, 7)` and `dispersion.cutoff=c(0.5, Inf)` for HVG selection; corrected batch effects for all 63 10x channels by calling `FindIntegrationAnchors` and `IntegrateData` with default parameters; ran `ScaleData` with default parameters and `RunPCA` with `seed.use=0` for PCA computation; ran `FindNeighbors` with `k.param=100`, `compute.SNN=FALSE`, `dims=1:50` and `nn.method="annoy"` to find k neighbors; ran `FindClusters` with `resolution=1.3`, `random.seed=0` and `algorithm="louvain"` or `algorithm="leiden"` to find clusters using either Louvain or Leiden algorithm; ran `RunTSNE` with `seed.use=0`, `dims=1:50` and `tsne.method="FIt-SNE"` to calculte tSNE-like visualization; ran `RunUMAP` with `seed.use=0`, `umap.method="umap-learn"`, `metric="correlation"`, `n.neighbors=15`, `min.dist=0.5` and `dims=1:50` to calculate UMAP-like visualization.

**SCANPY.** We first set `scanpy.settings.n_jobs=28`. Next, we ran `scanpy.pp.highly_variable_genes` with parameters `min_mean=0.0125`, `max_mean=7`, `min_disp=0.5`, `n_top_genes=2000` and `batch_key='Channel'` for HVG selection; imported `bbknn` python package directly and ran `bbknn.bbknn` with parameter `batch_key = 'Channel'` to correct batch effects; ran `scanpy.pp.scale` with parameter `max_value=10` and then `scanpy.tl.pca` with parameter `random_state=0` to compute PCA; ran `scanpy.pp.neighbors` with parameters `n_neighbors=100` and `random_state=0` to find k neighbors; ran `scanpy.tl.diffmap` with parameter `n_comps=100` to compute diffusion map; ran `scanpy.tl.louvain` and `scanpy.tl.leiden` with parameters `resolution=1.3` and `random_state=0` for Louvain and Leiden clusterings; ran `scanpy.tl.tsne` with parameters `use_rep='X_pca'`, `random_state=0` and `use_fast_tsne=True` to compute t-SNE; ran `scanpy.tl.umap` with parameter `random_state=0` to compute UMAP; ran `scanpy.tl.draw_graph` with parameters `random_state=0` and `iterations=500` to compute FLE.

**Pegasus.** We ran `pegasus cluster` with parameters `-p 28 --correct-batch-effect --knn-full-speed --diffmap --spectral-louvain --spectral-leiden --fitsne --net-umap --net-fle`.

#### 2.2 Function calls, commands and parameters used for benchmarking Pegasus, SCANPY and Seurat v3 on the 1.3 million mouse brain dataset (Table S3)

**Seurat.** Seurat failed to load this big dataset into memory.

**SCANPY.** SCANPY function calls and parameters were set exactly the same as in the bone marrow benchmarking experiment.

**Pegasus.** We ran `pegasus cluster` with parameters `-p 28 --correct-batch-effect --knn-full-speed --diffmap --spectral-louvain --spectral-leiden --fitsne --net-umap --net-fle --fle-memory 128`.

#### 2.3 Cloud benchmarking (Table 1)

For each of the three methods, we performed the following computational analyses: low quality cells/genes filtration, normalization to log expression space, HVG selection, batch correction, PCA, k-NN graph construction, clustering, tSNE-like and UMAP-like visualizations and find cluster-specific markers by differential expression analysis. For Cumulus, we additionally ran a putative cell-type annotation step.

**Seurat.** The `future` framework is used for parallelization. We first filtered low quality cells and converted counts to log expression levels as described in the paper (using Seurat functions). Next, we ran `FindVariableFeatures` with parameters `selection.method="vst"`, `mean.cutoff=c(0.0125,7)`, `dispersion.cutoff=c(0.5, Inf)` and `nfeatures=2000`. To correct batch effects, we ran `FindIntegrationAnchors` with parameter `dims = 1:20` and `IntegrateData` with parameter `dims = 1:20`. In addition, we used only 2 threads for batch correction. We have tried 10, 15, 20 and 32 threads and they all failed. We then ran `ScaleData` with default parameters and `RunPCA` with `seed.use=0` and `npcs=50` for PCA computation; ran `FindNeighbors` with `reduction="pca"`, `dims=1:50`, `nn.method='annoy'`, `k.param=100` and compute.SNN=TRUE to find k neighbors; ran FindClusters with resolution=1.3, random.seed=0 and algorithm="louvain" to compute clusters; ran RunTSNE with seed.use=0, dims=1:50 and tsne.method="FIt-SNE" to calculte tSNE-like visualization; ran RunUMAP with seed.use=0, umap.method="umap-learn", metric="correlation", n.neighbors=15 and dims=1:50 to calculate UMAP-like visualization. Lastly, we ran FindAllMarkers with parameters test.use="t" and random.seed=seed to find cluster-specific markers.

**SCANPY.** We first set scanpy.settings.n_jobs=os.cpu_count(). Then we filtered low quality cells and converted counts to log expression levels as described in the paper (using SCANPY functions). Next, we ran scanpy.pp.highly_variable_genes with parameters n_top_genes=2000 and batch_key='Channel' for HVG selection; ran scanpy.pp.scale with parameter max_value=10 and then scanpy.tl.pca with parameters n_comps=50, random_state=0 to compute PCA; imported bbknn python package directly and ran bbknn.bbknn with parameter batch_key = 'Channel' to correct batch effects; ran scanpy.tl.louvain with parameters resolution=1.3 and random_state=0 for Louvain clustering; ran scanpy.tl.tsne with parameters use_rep='X_pca', random_state=0 and use_fast_tsne=True to compute t-SNE; ran scanpy.tl.umap with parameter random_state=0 to compute UMAP. Finally, we ran scanpy.tl.rank_genes_groups with parameters groupby='louvain' and method='t-test' to find cluster-specific markers.

**Cumulus analysis step.** We ran the Cumulus WDL with the following parameters: zones="us-west1-b", num_cpu=32, memory=120G, correct_batch_effect=true, minimum_number_of_genes=0, knn_full_speed=true, run_louvain=false, run_spectral_louvain=true, run_net_umap=true, cluster_labels=spectral_louvain_labels, auc=false, fisher=false and annotate_cluster=true. In addition, since we did not compute diffusion maps, the spectral-Louvain was run on the principal component (PC) space.
